# Detection of hard and soft selective sweeps from *Drosophila melanogaster* population genomic data

**DOI:** 10.1101/2020.06.20.163261

**Authors:** Nandita Garud, Philipp W. Messer, Dmitri Petrov

## Abstract

Whether hard sweeps or soft sweeps dominate adaptation has been a matter of much debate. Recently, we developed haplotype homozygosity statistics that (i) can detect both hard and soft sweeps with similar power and (ii) can classify the detected sweeps as hard or soft. The application of our method to population genomic data from a natural population of *Drosophila melanogaster* (DGRP) allowed us to rediscover three known cases of adaptation at the loci *Ace*, *Cyp6g1*, and *CHKov1* known to be driven by soft sweeps, and detected additional candidate loci for recent and strong sweeps. Surprisingly, all of the top 50 candidates showed patterns much more consistent with soft rather than hard sweeps. Recently, Harris *et al.* 2018 criticized this work, suggesting that all the candidate loci detected by our haplotype statistics, including the positive controls, are unlikely to be sweeps at all and instead these haplotype patterns can be more easily explained by complex neutral demographic models. They also claim, confusingly, that these neutral non-sweeps are likely to be hard instead of soft sweeps. Here, we reanalyze the DGRP data using a range of complex admixture demographic models and reconfirm our original published results suggesting that the majority of recent and strong sweeps in *D. melanogaster* are first likely to be true sweeps, and second, that they do appear to be soft. Furthermore, we discuss ways to take this work forward given that the demographic models employed in such analyses are generally necessarily too simple to capture the full demographic complexity, while more realistic models are unlikely to be inferred correctly because they require fitting a very large number of free parameters.

## Introduction

Pervasive adaptation has been extensively documented in *Drosophila melanogaster*. Recent studies suggest that (i) ~50% of amino acid changing and non-coding substitutions in *D. melanogaster* evolution were adaptive, and (ii) there are abundant signatures of adaptation in the population genomic data detectable as reductions of neutral diversity in the regions of higher functional divergence and in the patterns of derived allele frequencies (*1*–*11*).

In three cases -- at the loci *CYP6g1*, *CHKov1*, and *Ace* – we specifically know the causal mutations and have functional hypotheses for the causes of adaptation (*12*–*18*). Intriguingly, in all three cases, there is strong evidence that adaptation was not driven by a single *de novo* adaptive mutation that rose to high frequency, but rather, multiple adaptive mutations. In the case of *Cyp6g1,* adaptive changes leading to resistance to DDT evolved via multiple insertions of *Accord* transposon in the 5’ regulatory region of the locus on different genomic backgrounds, as well as a duplication of the entire locus (*12*, *13*). At the *CHKov* locus, the adaptive change led to a higher resistance to organophosphates and viral infections and evolved by a transposon element insertion in the protein coding region of *CHKov*, which then segregated in the ancestral populations before rising to high frequency only recently (*14*, *15*). Finally, resistance to pesticides such as carbamates and organophosphates evolved via multiple independent point mutations at four highly conserved sites in the gene *Ace* on different genomic backgrounds on multiple continents (*16*–*18*). Not surprisingly, all three well-understood examples of adaptation show signatures of soft sweeps (**Figure 1**), in which multiple adaptive alleles rise to high frequency simultaneously at the same locus (*19*–*21*), and suggest that recent and strong adaptation is not mutation-limited in *D. melanogaster*.

**Figure 1:**
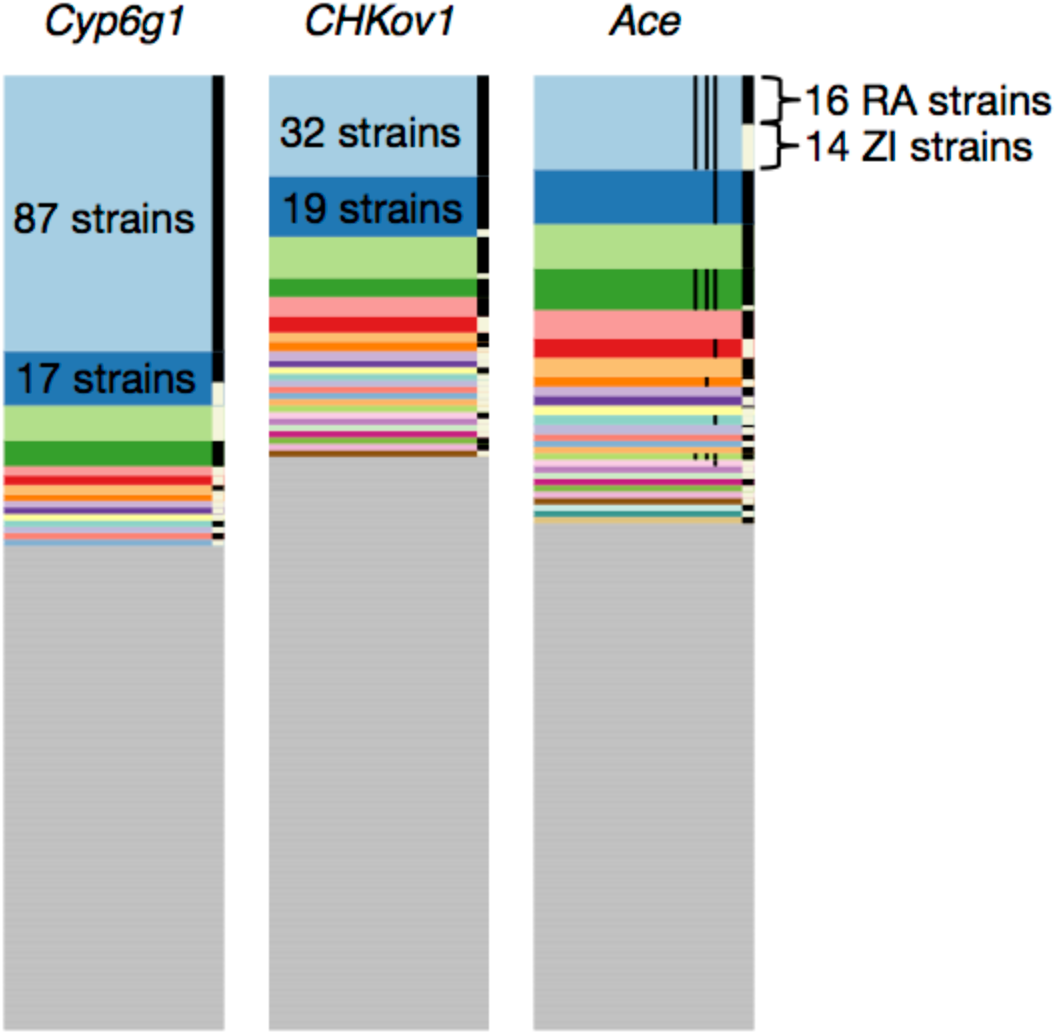
Haplotype frequency spectra at the *Cyp6g1*, *CHKov1*, and *Ace* loci. Recreated from Garud and Petrov 2016 (*17*). Haplotype frequency spectra at the three positive controls in a joint dataset, comprised of 300 Raleigh (RA) and Zambian (ZI) strains in 801 SNP windows, centered around the sites of the selective sweeps. 801 SNP windows in the joint data set correspond to slightly smaller analysis window sizes (<10 kb) in terms of base pairs on average than in the Raleigh or Zambian data alone. Each color bar represents a different, unique haplotype, and the height of the bar represents the number of chromosomes sharing the haplotype. The grey bars represent unique, singleton haplotypes in the sample. On the right side of each of the frequency spectra are black and white bars, indicating which strains are from RA and ZI, respectively. At all three positive controls, common haplotypes are shared across the two populations. The thin black lines shown in the haplotype spectrum for *Ace* correspond to the presence of three adaptive mutations that confer pesticide resistance.

These three empirical examples of soft sweeps at *Ace, CYP6g1,* and, *CHKov* were all defined experimentally and suggested that soft sweeps might be common. However, until recently, it was unknown how common soft selective sweeps are in the Drosophila genome. Most scans for detecting selective sweeps were specifically designed to detect signatures of unusually low diversity or the presence of a single common haplotype expressly associated with hard rather than soft selective sweeps (*2*, *22*–*29*), making it challenging to assess the frequency of soft sweeps (*30*–*33*).

To address this, we recently introduced novel haplotype homozygosity statistics for the detection and differentiation of hard and soft sweeps that are capable of detecting both hard and soft sweeps with similar power and then to determine whether detected sweeps are likely to be either hard or soft (*34*). Application of these statistics to the Drosophila Genetic Reference Panel (DGRP) (*35*), composed of 145 whole-genome sequences from a North Carolina *D. melanogaster* population, revealed several putative sweeps with unusually high haplotype homozygosity relative to expectations under several neutral demographic scenarios (**Figure 2**). The top 50 empirical outliers, which included the rediscovered positive controls at *CYP6G1, CHKov1,* and *Ace*, had multiple unusually long haplotypes present at high frequency, consistent with soft sweeps (**Figure 1**). We found that simulations of hard sweeps were unable to produce signatures observed in the data, whereas simulations of soft sweeps easily could. Subsequent studies found that soft sweeps seem to be common not only in this North American population, but also in Sub Saharan populations of *D. melanogaster* (*17*, *18*, *36*). Finally, these haplotype homozygosity statistics have been applied to several other organisms including pigs (*37*, *38*), dogs (*37*), cattle (*39*, *40*), soy beans (*41*), and humans (*42*) to identify hard and soft sweeps, and have become standard summary statistics in machine learning methods for detecting selection (*43*, *44*).

**Figure 2:**
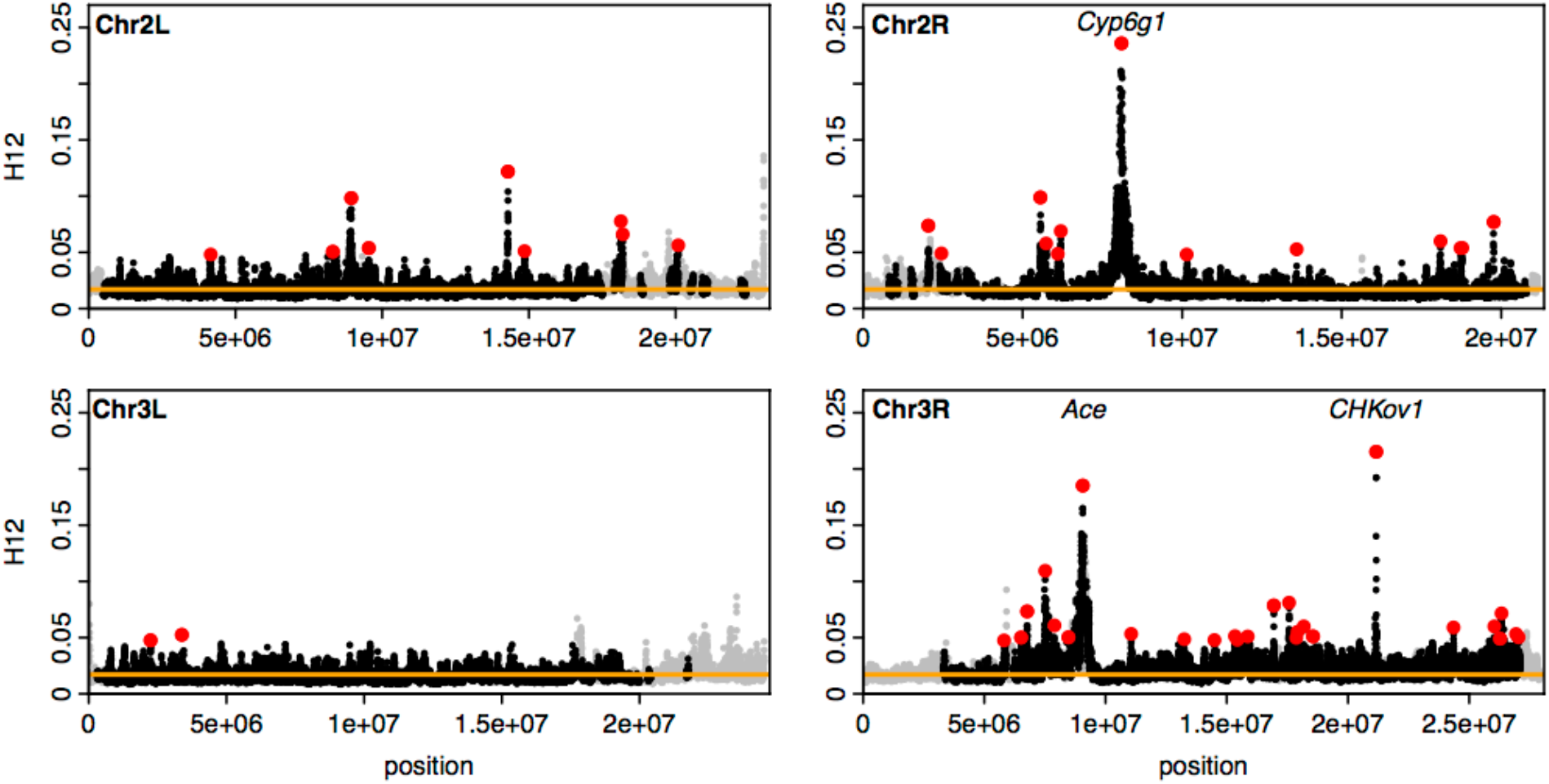
H12 scan of DGRP data. Recreated from Garud *et al.* 2015 (*34*). Scan of the four autosomes using the H12 statistic. Each point indicates an H12 value computed in a 401 SNP window. Grey points indicate regions excluded from the analysis with recombination rates lower than 5×10^-7 cM/bp. The orange line represents the 1-per-genome FDR line calculated from simulations of a neutral model with constant population size of 10^6. Red points indicate the top 50 extreme outlier peaks relative to the 1-per-genome FDR line. Three positive controls are indicated at *Ace, Cyp6g1,* and *CHKov1.*

Harris et al. 2018 (*45*) recently reanalyzed the DGRP data using our statistics and argued that that there is scant evidence for abundant recent strong selective sweeps in the North American *D. melanogaster* population. They claim that correct neutral admixture models naturally generate detected haplotype signatures in the absence of positive selection and thus most of the detected signatures do not correspond to selective sweeps. They also argue that if these sweeps exist, then they would be hard rather than soft sweeps. Here we re-evaluate our analysis using a range of demographic models and show that our previous findings stand. We then discuss the reasons for the different conclusions from Harris et al. 2018 (*45*), implications of these results, and directions for future work.

## Results

### Summary of our previous results in Garud et al. 2015

The increase in frequency of an adaptive allele is expected to also lead to the increase in frequency of the linked haplotype (*46*–*48*). Such an increase of haplotype frequency is expected to elevate levels of haplotype homozygosity (H1) in the vicinity of the selected locus (*25*, *26*, *28*, *29*, *49*). H1 is defined as

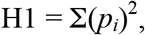

where *p*_*i*_ is the frequency of the i^th^ most common haplotype. H1 is expected to be elevated for both hard and soft selective sweeps, but hard sweeps should still have higher H1 values than soft sweeps, given that soft sweeps bring multiple haplotypes to high frequency.

In Garud *et al.* 2015 (*34*) we define a similar haplotype homozygosity statistic H12, which combines the frequencies of the first and second most common haplotypes into a single frequency and is defined as follows:

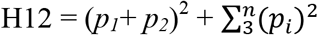

Using extensive simulations, we showed that H12 has reasonably similar power to detect hard and soft sweeps with a slight bias in favor of hard sweeps (*34*).

The application of these statistics requires one to define a window size. Longer windows should have lower false positive rates for distinguishing selection from neutrality, but simultaneously have reduced ability to detect weaker sweeps. In Garud *et al.* 2015 (*34*) we used windows of 401 SNPs in length (~10Kb in Drosophila), and we show that this biases our analysis towards detecting sweeps with selection coefficients (*s*) >= 0.1%. Note that most of the detected sweeps span multiple analysis windows with the peaks ranging from ~11kb to ~870 kb, and with half of the peaks over 100kb (**Figure 2**), suggesting that the identified sweeps were substantially stronger on average than *s* = 0.1%.

We tested a range of neutral models fitting overall polymorphism levels in the data and found that these rarely generate elevated values of H12 on such long length scales (*34*). We specifically considered six models of increasing complexity **(Figure 3)**. We included four simple models that were fit to site frequency spectrum-based summary statistics measured from short introns in the DGRP data: two constant population size models (the standard *Ne*=10^6^ and *Ne*=2.2*10^6^ which fit the levels of diversity better) and two bottleneck models with varying bottleneck durations and sizes. Finally, we included two complex admixture models inferred by Duchen et al. 2013 (*50*) using an approximate Bayesian computation (ABC) approach and data from 242 short intronic and intergenic fragments from the X-chromosome. These models were fit to both site frequency spectrum-based summary statistics (number of segregating sites, S/bp, and average nucleotide diversity, Pi/bp) and LD measured on short length-scales (~500bp) using Kelly’s ZnS. Using their ABC method, Duchen et al. 2013 (*50*) inferred a posterior distribution for each of the 11 parameters for the admixture models.

**Figure 3:**
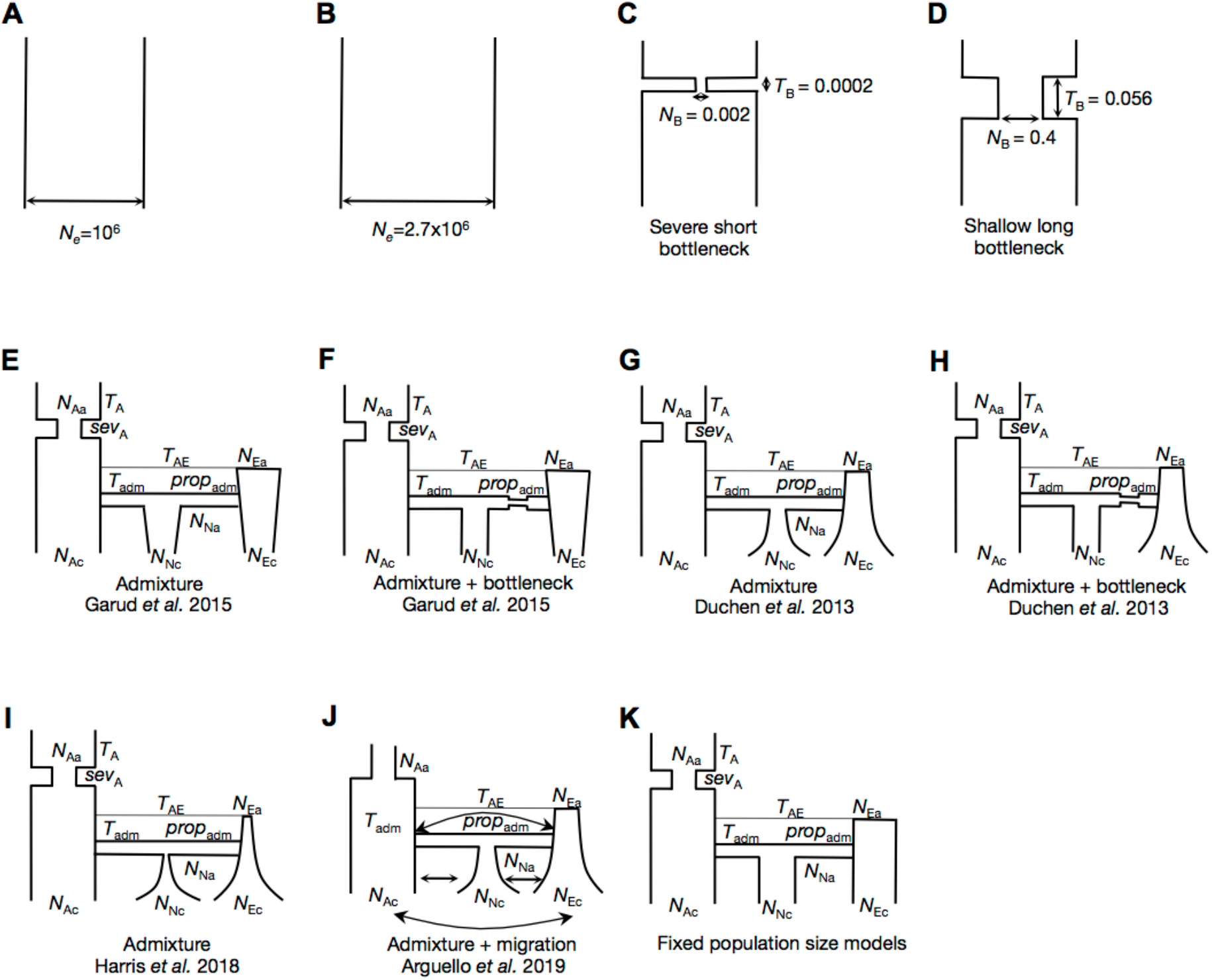
Neutral demographic models. Diversity statistics were measured in simulations of 11 neutral demographic models: (A) A constant *N*_*e*_ = 10^6^ model (B) A constant *N*_*e*_ = 2.7×10^6^ model (fit to Watterson’s *θ*_W_ measured in autosomal short introns in DGRP data) (C) A severe short bottleneck model fit to Pi and S in autosomal short introns in DGRP data (D) A shallow long bottleneck model fit to Pi and S in autosomal short introns in DGRP data (E) The implemented admixture model in Garud *et al.* 2015 (F) The implemented admixture + bottleneck model in Garud *et al.* 2015 (G) The admixture model proposed by Duchen *et al.* 2013 (H) The admixture + bottleneck model proposed by Duchen *et al.* 2013 (I) The implemented admixture model in Harris *et al*. 2018 (J) The admixture model proposed by Arguello *et al.* 2019 (K) A variant of the Duchen *et al.* 2013 admixture model where North America, Europe, and Africa have fixed population sizes.

Upon revisiting the Duchen et al. 2013 (*50*) models for the present paper, we found that the model implemented in Garud et al. 2015 (*34*) was a variation of the model published in Duchen et al 2013 (*50*). Instead of an expansion in North America and Europe, we had implemented roughly constant population sizes in North America and Europe (**Figures 3E** **and** **3F**). Despite this difference from the published model, all six implemented models fit the autosomal DGRP data in terms of S/bp, Pi/bp and decay in short-range LD (**Figure 4**). Both H12 and long-range LD (~10Kb) in simulated models were depressed compared to the observed data (**Figures 4C** **and** **5**). Specifically, the data showed much slower decay of LD on the scale of 10kb and substantially larger median haplotype homozygosity (H12). The distribution of H12 in the data also had a much longer tail compared to simulations (**Figures 5** **and** **S1**). Indeed, the observed median values of H12 in the data are similar to levels expected only once in the genome under these neutral models. We interpreted the elevation of long-range LD and haplotype homozygosity in long windows in the data as evidence of positive selection.

**Figure 4:**
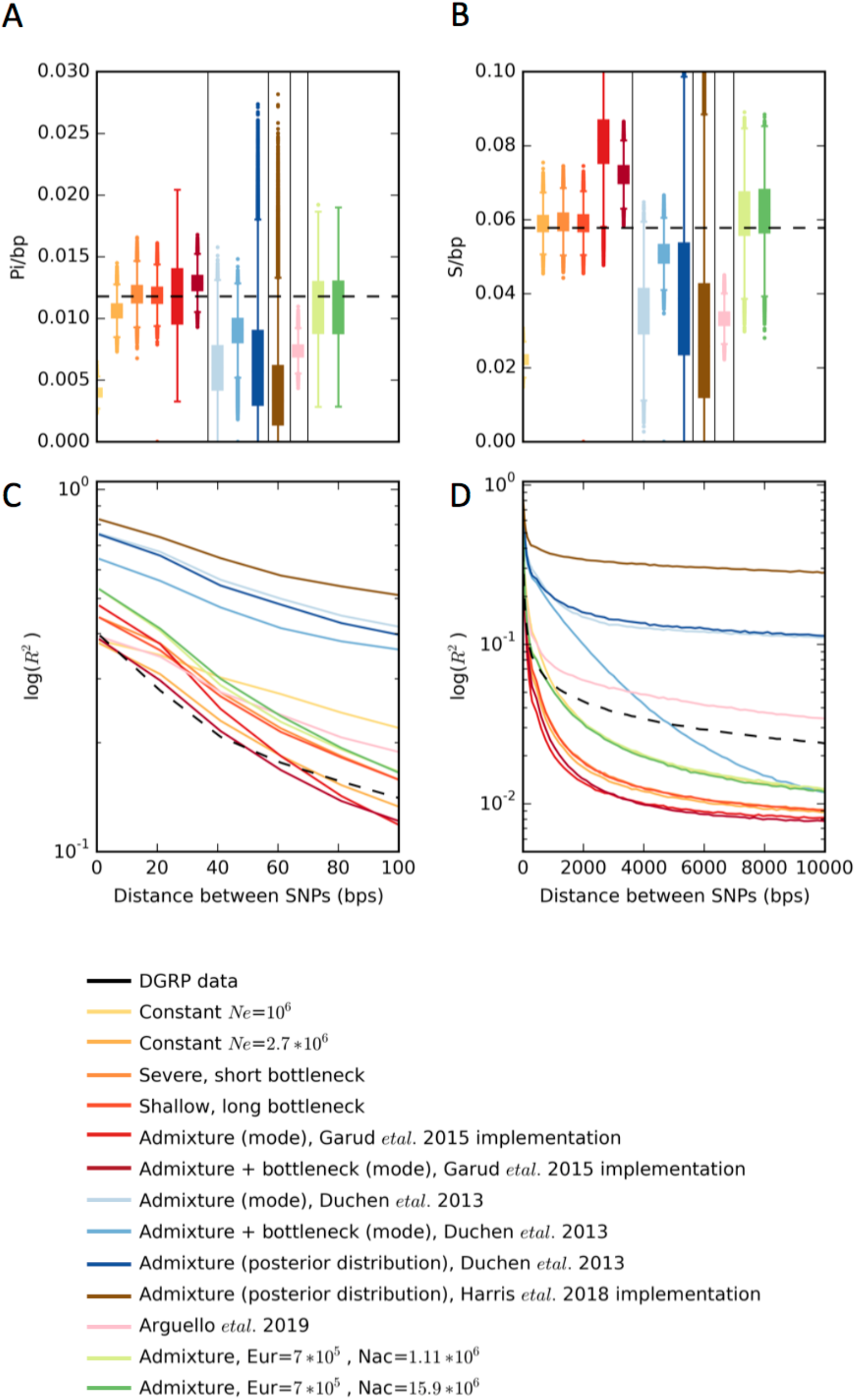
Distributions of Pi, S, and linkage disequilibrium in data and simulations. Distributions of (A) Pi/bp, (B) S/bp, (C) short range LD (R^2^), and (D) long range LD measured in DGRP data and simulated neutral demographic models. Simulations were generated with a recombination rate *ρ* = 5×10^−7^ cM/bp. Diversity statistics were calculated in DGRP data in genomic regions with *ρ* ≥ 5×10^−7^ cM/bp. The horizontal dashed lines in (A) and (B) depict the median Pi/bp, S/bp, and H12 values measured in DGRP data. For each model, statistics from 1.3×10^5^ simulations are plotted in (A) and (B). The dashed black lines in (C) and (D) correspond to mean LD values computed in DGRP data. LD in simulations was estimated from 1×10^7^ pairs of SNPs.

**Figure 5:**
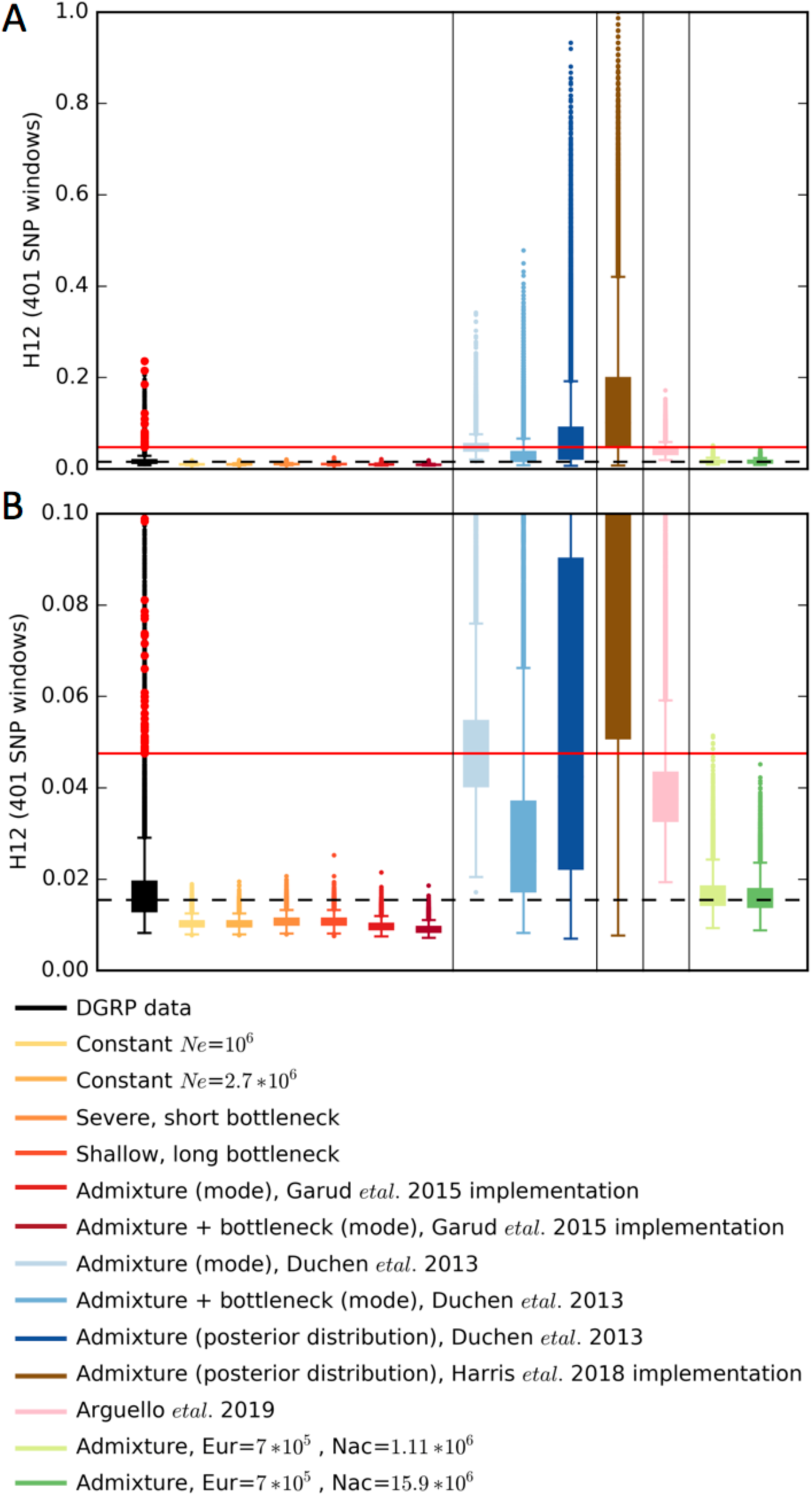
H12 distributions in data and simulations. Distributions of H12 in 401 SNP windows. Shown are the (A) full distribution and (B) truncated y-axis for visual clarity. Simulations were generated with a recombination rate *ρ* = 5×10^−7^ cM/bp and H12 was calculated in DGRP data in genomic regions with *ρ* ≥ 5×10^−7^ cM/bp. The horizontal dashed line indicates the median H12 value in DGRP data and the horizontal red line indicates the lowest H12 value for the top 50 peaks. H12 values from 1.3×10^5^ simulations for each model are plotted. The distribution of genome-wide H12 values measured in DGRP data is shown in black. Overlaid in red points are the H12 values corresponding to the top 50 empirical outliers in the DGRP scan.

The lack of the model fit with the bulk of the H12 values in the data presents a problem for identifying selective sweeps with elevated homozygosity. Thus, we elected to be conservative. First, we defined a 1-per-genome false discovery rate for each demographic model by performing 1.3×10^5 simulations (>10 times the number of analysis windows observed in the data) for each model. The FDRs, corresponding to the 10^th^ highest H12 value in the distributions, were approximately equal to the median H12 value observed in the data. Several genomic regions had ‘peaks’ of elevated H12 values that were especially unlikely to be generated by neutrality. These regions corresponded to our candidate selective sweeps (**Figure 2**). We then focused on the 50 empirical outliers to further characterize as hard or soft. These peaks were defined by identifying the window with the highest H12 value and finding all consecutive windows in both directions with H12 values exceeding the 1-per-genome FDR. These candidates all had maximum H12 at least eleven standard deviations away from the median H12 value in the data after fitting a Gaussian distribution to the bulk of the data **(Figure S4)** (Methods). The top 3 outliers were the positive controls, *Ace, Cyp6g1, CHKov1,* confirming that H12 has the ability to detect known soft selective sweeps that arose from multiple *de novo* mutations or standing genetic variation.

### Can admixture generate elevated haplotype homozygosity?

Harris et al. (*45*) claim that the admixture model proposed by Duchen et al. 2013 (*50*) can easily generate all the elevated H12 values in the data, suggesting that the selective sweeps identified by H12 are false positives.

Given that our original implementation of the admixture model in Garud et al. 2015 (*34*) was a variant of the Duchen et al. 2013 (*50*) model, we tested Harris et al.’s (*45*) claim by implementing the model specified in their supplement, which also differs from the Duchen et al. 2013 (*50*) model (methods). To more broadly consider the Harris et al.’s (*45*) claim that appropriate demographic models can easily generate the distribution of H12 values observed in the data, we also tested the Duchen et al. 2013 (*50*) model both with the mode of the posterior distributions of the 11 parameters for the admixture model and by drawing parameters from the 95 CIs of the posterior distributions, as was done in Harris et al. 2018 (*45*). Note that Harris et al. 2018 (*45*) did not use a joint posterior distribution, leading to the possibility that many of the parameter combinations may not correspond to realistic scenarios. We also tested a variant of the admixture model proposed by Duchen et al. 2013 (*50*) that included a bottleneck in the founding European population (**Figure 3H**). Finally, we also tested a new admixture model with migration between Africa, Europe, and North America (**Figure 3J**) recently proposed by Arguello et al 2019 (*51*).

The Duchen et al. 2013 (*50*) model and Harris et al. 2018 (*45*) implementation of the model generate S/bp and Pi/bp values that are 2-fold lower than the median values measured from short introns in the DGRP data (**Figures 4A** **and** **4B**). More strikingly, however, H12 is extremely elevated compared to values observed in the DGRP data (**Figures 5** **and** **S2**), e.g the bulk of the distribution of values generated in simulations is non-overlapping with the bulk of the distribution of values from genome-wide data. This elevation is likely due to the sharp bottlenecks specified in the models, especially in the Harris et al. 2018 implementation where the bottleneck size is 4 times smaller than reported by Duchen et al. 2013 (*50*). The elevation is even more pronounced when drawing parameters from the 95 CIs. In some cases, H12 almost approaches 1, implying that these models predict essentially no variation in the DGRP in the large ~10kb window sizes used for our analysis. Consistent with elevated haplotype homozygosity, the Duchen et al. 2013 (*50*) models produced elevated pairwise LD compared to observations in the data (**Figures 4C** **and** **4D**). The mismatch between the expected values of all the statistics in the Arguello et al. 2019 (*51*) model and the data is less pronounced compared to the Duchen et al. 2013 (*50*) model, presumably because migration replenishes some of the diversity lost during the extreme bottlenecks, but nevertheless there is a significant mismatch.

At first glance, the elevated haplotype homozygosity produced by the Duchen et al. 2013 (*50*) model might suggest that the peaks observed in the DGRP data (**Figure 2**) could be explained by the admixture model. However, the S/bp, Pi/bp, H12, and LD values produced by this admixture model deviate significantly from genome-wide summary statistics in the data. In particular, the distribution of H12 values in the data has a very specific distribution that the simulations of neutrality cannot match (**Figures 5** **and** **S1**–**3**). Almost 80% of the analysis windows in the DGRP data have H12 values within 2 standard deviations from the median, after fitting a Gaussian to the bulk of the distribution (**Methods**, **Figure S4**). This is followed by a long tail of H12 values that includes the values for the top 50 peaks, which are >= 11 standard deviations away from the median, indicating that these peaks are indeed genome-wide outliers (**Figure S4**). By contrast, the bulk of the distribution generated by the Duchen et al. 2013 (*50*) admixture model surpasses the median and bulk of the distribution of H12 values in the data (**Figure 5**). This lack of fit of the admixture model to the data is problematic for the inference of selective sweeps: if the tail of the distribution of H12 values from data can be explained by neutrality, then the bulk of the distribution should also be explainable by neutrality. These admixture models do not recapitulate the distribution observed in the data, and instead produce extremely high levels of homozygosity that are incompatible with the data.

One reason for the lack of fit of H12 measured in the data and the simulated admixture model could be that Duchen et al. 2013 (*50*) initially fit the model to the 242 X-chr fragments of ~500bps using SFS statistics and short-range LD statistics (Kelly ZnS), whereas we analyzed autosomal data. Although Duchen et al. 2013 (*50*) showed that the model extrapolated to autosomes by fitting to ~50 intronic and intergenic regions on the 3^rd^ chromosome, the models do not fit diversity patterns on short introns, which are putatively the most neutral part of the genome (*52*). Additionally, Duchen et al. 2013 (*50*) did not require that long-range haplotype homozygosity on the scale of ~10kb fit the data, which is the main source of discrepancy between the models and the data.

Models that fail to recapitulate the bulk of the diversity statistics from the data are unlikely to accurately capture the true demographic history of the population. These models are not appropriate for inferring sweeps because they do not set a realistic baseline for the expected diversity pattern in a neutral scenario.

### Inference of new demographic models

In this section, we test whether there are variants of the Duchen et al. 2013 (*50*) and Arguello et al. 2019 (*51*) admixture models that can achieve a better fit with regard to multiple relevant summary statistics in the DGRP data. The models we explore are not intended to be a true representation of the demography of the North American population of Drosophila. Instead, our goal is to see whether we can find an admixture model that reasonably fits both SFS and LD-based genome-wide statistics in the data and can also generate tails of elevated H12 values that may explain the outlier peaks observed in the data. We do not claim that other models cannot fit the data equally well or better.

We tested four classes of variants of the Duchen et al. 2013 (*50*) and Arguello et al. 2019 (*51*) admixture models (**Figure S5**). First, we tested models with constant population sizes in North America and Europe (**Figures 3K** and **S5A**), because the Garud et al. 2015 (*34*) implementation (**Figure 3E**), which had effectively constant population sizes for these two populations, fit the data well in terms of S/bp and Pi/bp. Second, we tested models with varying amounts of growth in North America and Europe (**Figure S5B**). Third, we tested models with varying proportions of admixture (**Figure S5C**), and fourth, we tested models with varying amounts of migration between the continents (**Figure S5D**). For each of these models, we held almost all parameters constant at the mode of the posterior distributions inferred by Duchen et al. 2013 (*50*). The only parameters we varied were those relevant to the model being tested (e.g. proportion of admixture, amount of migration, or rate of growth). Where applicable, the values of these parameters were chosen to span the ranges of the 95CI inferred by Duchen et al. 2013 (*50*). These variable parameters are highlighted in red in **Figure S5**. In sum, we tested a total of 74 admixture model variants. Supplemental **Figures S6**–**18** show the distributions of summary statistics S, Pi, H12, short-range LD and long-range LD generated by these models.

The majority of the models tested do not fit the data well, whereby the median values of S/bp, Pi/bp, and H12 measured in the data lie outside the 25^th^ and 75^th^ quantiles measured from simulations (**Figures 4**, **5**, **and** **S1**–**3**). Models that produce extremely high H12 values and low S and Pi values generally have small founding population sizes. Models with depressed H12 values and elevated S and Pi values have larger founding population sizes. Many models fit some summary statistics reasonably well, but no single model fits all five summary statistics.

Only 3 of the 74 models we tested generate distributions overlapping the median genome-wide S, Pi, and H12 values (**Figures S3** **and** **S8**). These models have constant population sizes in North America and Europe (**Figures 3K** and **S5A**) of magnitudes similar to the one implemented in Garud et al. 2015 (*34*). Specifically, the well-fitting models have large North American population sizes (>= 1.11*10^6), and intermediate European population sizes (~0.7*10^6). The distributions of S, Pi, H12, and LD for two of these models are shown in **Figure 4** as a comparison with all other models considered in this paper so far.

While the three well-fitting models generate S, Pi, and H12 values that overlap the median values measured from genome-wide data, they cannot generate long tails of elevated H12 values. The 1-per-genome FDR values observed in simulations for these models have H12 values that are lower than even the 50^th^ ranking peak in the DGRP H12 scan. This suggests that given a reasonably well-fitting model, the top 50 H12 peaks observed in the DGRP data are still outliers under any of the current models.

### Distinguishing hard versus soft sweeps with the H2/H1 statistic

In Garud et al. 2015 (*34*), we analyzed whether the haplotype patterns observed among the top 50 peaks are more consistent with hard or soft sweeps. To do so, we introduced a second haplotype homozygosity statistic, H2/H1, to distinguish hard from soft sweeps.

H2 is haplotype homozygosity computed excluding the most common haplotype:

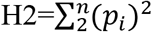

H2 is expected to be small for hard sweeps because the main contributing haplotype to homozygosity is excluded. However, it is expected to be larger for soft sweeps since there should be multiple adaptive haplotypes at high frequency. H2/H1 augments our ability to distinguish hard and soft sweeps since it is even smaller for hard sweeps and larger for soft sweeps than H2.

While hard sweeps and neutrality cannot easily generate both high H12 and H2/H1 values, soft sweeps can. Hence, the H2/H1 statistic is powerful for discriminating hard and soft sweeps only when applied to candidate selective sweeps with H12 values exceeding expectations under neutrality. Additionally, H2/H1 is inversely correlated with H12 values (*53*). Thus, H12 and H2/H1 must be jointly applied when H12 is sufficiently high to make inferences about the softness of a sweep.

In Garud et al. 2015 (*34*), we tested whether the H12 and H2/H1 values for the top 50 peaks are more consistent with hard versus soft sweeps. Specifically, we categorized sweeps as hard versus soft by computing Bayes factors: BF = P(H12obs, H2obs/H1obs | soft sweep)/P(H12obs,H2obs/H1obs | hard sweep), whereby H12obs and H2obs/H1obs were computed from data, and hard and soft sweeps were simulated by drawing partial frequencies, selection strengths, and ages from uniform prior distributions (Methods). By using a Bayesian approach, we can then integrate over a wide range of evolutionary scenarios instead of testing a single point hypothesis. We found in our application of H2/H1 and H12 to the DGRP data under both constant Ne models and our implementation of the admixture models that the top 50 peaks have H12 and H2/H1 values more consistent with soft sweeps than hard sweeps.

We repeated the BF analysis (Methods) with the new admixture models (**Figure 3K**) inferred in this paper and found that the majority of the sweeps are soft and ~3-4 are hard depending on the model under consideration (**Figure 6**). Visual inspection of the haplotype frequency spectra for the top 50 peaks confirms that multiple haplotypes are present at high frequency (**Figure 7**) for most peaks.

**Figure 6:**
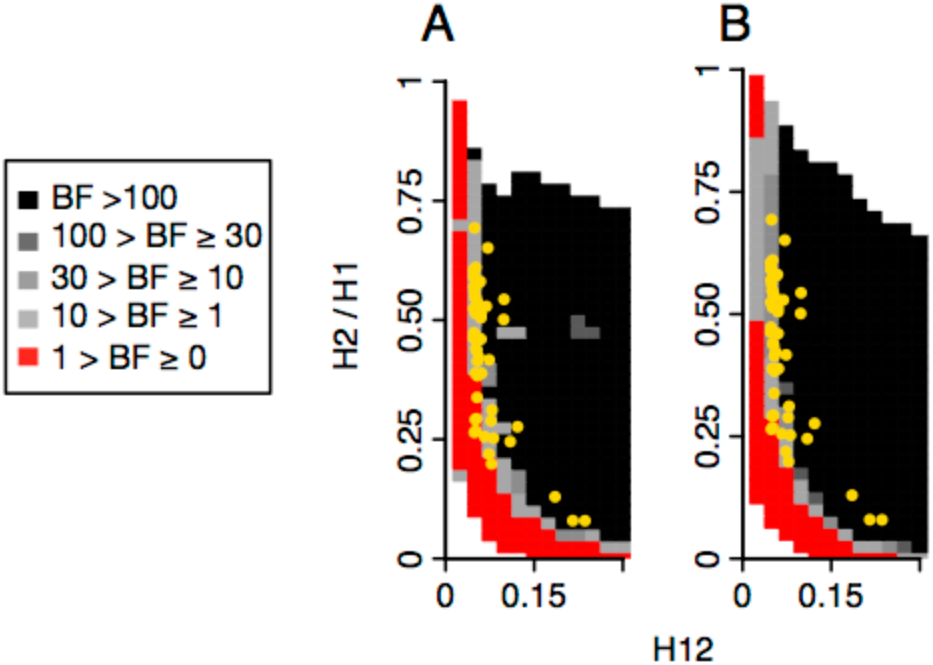
Range of H12 and H2/H1 values expected for hard and soft sweeps under two admixture models. Bayes factors (BFs) were calculated for a grid of H12 and H2/H1 values to demonstrate the range of H12 and H2/H1 values expected under hard versus soft sweeps. Panels A and B show results for variations of the admixture model proposed by Duchen *et al.* 2013, where Africa, North America, and Europe have constant population sizes. In (A), the population sizes for North America and Europe were held constant at 1,110,000 and 700,000 individuals, respectively. In (B), the population sizes for North America and Europe were held fixed at 15,984,500 and 700,000 individuals, respectively. BFs were calculated by computing the ratio of the number of soft sweep versus hard sweep simulations that were within a Euclidean distance of 10% of a given pair of H12 and H2/H1 values. Red portions of the grid represent H12 and H2/H1 values that are more easily generated by hard sweeps, while grey portions represent regions of space more easily generated under soft sweeps. Each panel presents the results from 10^5^ hard and soft sweep simulations, respectively. Hard sweeps were generated with *θ*_A_ = 0.01 and soft sweeps were generated with *θ*_A_ = 10. A recombination rate of *ρ* = 5×10^−7^ cM/bp was used for all simulations. The H12 and H2/H1 values for the top 50 empirical outliers in the DGRP scan are overlaid in yellow.

**Figure 7:**
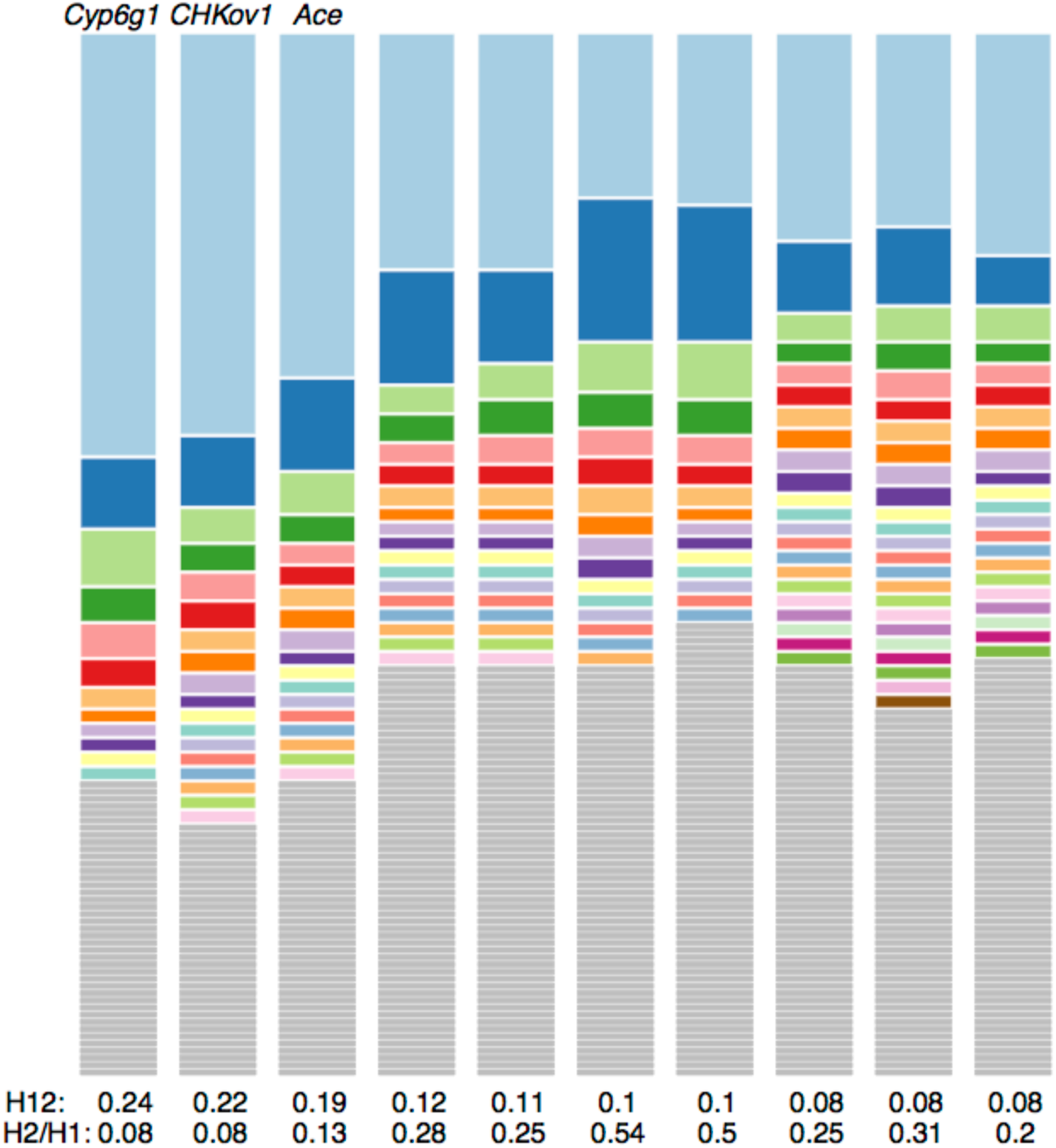
Haplotype frequency spectra for the top 10 peaks in the DGRP data. Haplotype frequency spectra for the top 10 peaks in the DGRP scan. For each peak, the frequency spectrum corresponding to the analysis window with the highest H12 value is plotted.

To make their argument that H2/H1 does not have power to distinguish hard and soft sweeps, Harris et al. 2018 (*45*) assessed whether the top 50 peaks have H2/H1 values consistent with hard versus soft sweeps (even though they also claimed that that these peaks are not sweeps to begin with). In their Figure 1D, they conclude that H2/H1 does not have discriminatory power. However, the H2/H1 values for the top 50 peaks lie within the bulk of the distribution generated by soft sweeps and in the tail of the distribution generated by hard sweeps (see their Figure 1D), which appears at odds with their conclusion. Despite their claims that H12 and H2/H1 lack discriminatory power, Harris et al. 2018 (*45*) also computed Bayes factors (BF) in their Figure S1 and showed that after correctly conditioning on matching H12 and H2/H1 values for the top 50 peaks, the majority of the peaks have values that are consistent with soft sweeps. Thus, Harris et al. 2018 (*45*) obtain the same result as in Garud et al. 2015 (*34*).

## Discussion

Whether hard versus soft sweeps are common is a topic of much debate. While multiple empirical studies have revealed evidence for soft sweeps in a wide range of organisms including *D. melanogaster, P. falciparum* (*54*, *55*), viruses (*56*), humans (*42*, *57*–*60*), dogs (*37*), amongst others (*21*), several articles claim that there is unfounded enthusiasm for soft sweeps and that in fact, they are not as pervasive as the evidence suggests (*45*, *61*, *62*). Specifically, Harris et al. 2018 (*45*) suggest that the claim in Garud et al. 2015 (*34*) that there is abundant evidence for many strong and recent soft sweeps in the *D. melanogaster* populations is not supported after appropriate applications of demographic models and generating appropriate null distributions. Here we carry out a range of additional analyses and reassert the claims of Garud et al. 2015 (*34*).

In Garud et al. 2015 (*34*), we developed the haplotype homozygosity statistics H12 and H2/H1 to systematically detect and differentiate hard and soft sweeps from population genomic data. Our application of these statistics to the DGRP data from North Carolina revealed that soft sweeps are common in this population. Among the top candidates in our scan were the positive controls at *Ace*, *Cyp6g1*, and *CHKov1*. Corroborating our results, we found that approximately half of our sweep candidates were also identified with the popular statistic, *iHS* (*28*), which was designed to detect partial hard sweeps. Finally, we found that soft sweeps are common in a Zambian population as well (*17*), suggesting that any particular demographic history for a given population is not driving the signal of multiple haplotypes at high frequency. Independently, (*44*) found that soft sweeps are prevalent in this same population of Zambia.

To ensure low false positive rates, we excluded related individuals and tested for substructure. Additionally, we utilized large analysis window sizes of 401 SNPs (corresponding to ~10kb), since haplotypes of such length are not expected to be at high frequency by chance. Note that we use windows of constant number of SNPs to avoid the issue of H12 co-varying with the number of SNPs in the window. Windows defined in terms of the number of SNPs automatically extends the physical lengths of the windows in regions of low diversity. Such longer windows should then have proportionately higher recombination rates reducing the expected H12, and thus reducing the probability of false positives. In practice, we also eliminated regions of low recombination (rho <5*10^−7 cM/bp) from the data, as regions of low recombination rate can show elevated false-positive rates in haplotype-bases tests of selection (*63*). By contrast, Harris et al. 2018 (*45*) chose to perform their analyses in 10kb windows, although we note that it is unclear how the H12 and H2/H1 values plotted in their Figure 1 were identified -- plausibly they correspond to the 50 identified peaks in Garud et al. 2015 (*34*). It is unclear why the plotted values appear to correspond to the values in Garud et al. 2015 (*34*) despite the application of different approaches. It is also unclear whether windows of particularly low nucleotide diversity had been eliminated from the Harris et al.’s 2018 (*45*) scan or whether any scan was performed at all.

In Garud et al. 2015 (*34*), we tested several demographic models and found that while they do match Pi and S, they tend to generate values of H12 that are lower than in the data (**Figures 4** **and** **5**). While we remained agnostic to the cause of this inflation of H12 in the data (misspecification of demography or pervasive draft or both), we chose to focus on empirical outliers as a conservative approach. Our belief was that in general it is not yet possible to ensure that any demographic model is correct and thus the focus on empirical distributions is warranted.

Harris et al. 2018 (*45*) claim that reasonable demographic models fit the data well, but upon closer inspection of the models they tested, we find that they do not. Specifically, the Duchen et al. 2013 (*50*) model generates values of S and Pi that are 2-fold lower than the median values in the data, and extremely elevated H12 values that approach 1, suggesting that the North Carolina population should be almost monomorphic while in fact the DGRP data does not have such extreme H12 values. The lack of a well-fitting demographic model is problematic for the inference of selective sweeps. In this case, is the model plausible given that such high homozygosity is never even observed in the data? If the bulk of H12 values in this model are much higher compared to genome-wide levels of H12, then it is impossible to ascertain whether localized regions of high homozygosity in the data are significant departures from neutral expectations. Recently, Arguello et al. 2019 (*51*) inferred a new admixture model for North American Drosophila, which includes migration between Africa, Europe, and North America. However, this model does not fit H12, S, and Pi in the DGRP data either.

Thus, in the absence of a well-fitting null, we were inspired to look for a more reasonable null model to determine if such a model could in fact generate long H12 tails. We tested over 70 different versions of the admixture models and found that the majority of the tested models did not fit the data (**Figures S6–18**). When the founding population sizes of Europe and North America were very small, the models predicted a sharp depression of diversity. When the founding population sizes were too large, the models predicted very high diversity unobserved in the data. The models that did fit the data reasonably well in terms of S, Pi, and H12 were the ones with fixed population sizes in North America and Europe, similar to the one implemented in Garud et al. 2015 (*34*). We emphasize that these inferred models are not intended to be the ‘correct’ models, especially since LD measured with R^2 still does not match the data. Rather, they are useful for ascertaining whether a model that can fit multiple summary statistics in the data can also generate a long tail of H12 values. Future work that exhaustively searches the parameter space for a model that fits all summary statistics is needed.

The 50 peaks that we identified in Garud et al. 2015 (*34*) are all in the extreme tails of the new models that fit Pi, S, and now the bulk of the H12 distribution. These 50 peaks have H12 values that are more than 11 standard deviations away from median H12 value in the data (**Figure S4**), providing additional evidence that these peaks are outliers given a normal distribution that fits the bulk of the data quite well.

Detecting selective sweeps is only the first goal. The next goal is to distinguish hard and soft sweeps from each other. For this purpose, Garud et al. 2015 (*34*) developed the statistic H2/H1. Harris et al. 2018 (*45*) claim that H2/H1 cannot distinguish hard and soft sweeps, even though in their implementation they found that the observed H2/H1 values are in the tail of the values generated by hard sweeps and firmly within the bulk of the distribution generated by soft sweeps.

An additional reason for Harris et al.’s (*45*) conclusion that H2/H1 does not have sufficient power to distinguish hard versus soft sweeps is that they did not correctly condition on both H12 and H2/H1 in their Figure 1. H2/H1 being high alone is insufficient to determine if a sweep is soft because non-sweeps can easily generate high H2/H1 values. When H12 values are high and we do have evidence of a sweep, H2/H1 does in fact has high power to distinguish hard and soft sweeps from each other (*34*, *53*). Conditioning on the highest H12 value in a peak can also avoid confounding issues like soft shoulders (*61*), in which a hard sweep decays due to recombination and mutation events and results in soft sweep-like patterns a short distance away from the sweep center. Indeed, the H12 and H2/H1 statistics have become important discriminating statistics for several recent machine learning methods that detect and differentiate hard and soft sweeps (*36*, *42*, *43*). Moreover, Figures 1D and S1 from Harris *et al.* 2018 (*45*) show that after conditioning on H12 values, the top 50 peaks’ H12 and H2/H1 values are consistent with soft sweeps and not hard sweeps, even if they did not state it.

Regardless of the exact statistical methodology used or underlying demographic model, it is a fact that soft sweeps do occur. In *D. melanogaster* alone, there are three well-documented examples of soft sweeps at *Ace*, *Cyp6g1*, and *CHKov1* using direct observations of the same allele on distinct genomic backgrounds (*12*–*18*). More broadly, soft sweeps have been abundantly documented using a variety of methods, data sets and organisms (*21*). Our work here is not the final word on the topic as future statistical developments may enable us to better quantify rapid adaptation from population genomic data. Note that despite having tested more than 70 models, none could fit every summary statistic in the data. Thus, it is important to acknowledge that there may not be any purely neutral model that can explain the diversity patterns observed in the data. Factors such as linked selection (*64*), background selection (*65*–*67*), seasonal adaptation (*68*), local adaptation (*69*), and variable recombination rates (*70*) could all be contributing to diversity patterns in the data. Thus, a combination of demographic and selection forces may be needed to be jointly inferred to be able to fully match diversity patterns in the data. Identifying statistics capable of detecting selection that are robust to the misspecification of demographic and selective models might be one profitable direction for future research given how complex and strong evolutionary forces are known to be.

## Methods

### Simulations of neutrality and selection

Neutral simulations were generated with the coalescent simulator MS (*71*), and selection simulations were generated with MSMS (*72*). All samples consisted of 145 chromosomes to match the sample depth of the DGRP data analyzed in Garud et al. 2015 (*34*). Simulations were generated with a neutral mutation rate of 10^−9^ events/bp/gen (*73*) and a recombination rate of 5*10^−7 cM/bp.

To simulate hard and soft selective sweeps, we varied the adaptive mutation rate, *θ*_A_ = 4*Ne*mu_A. Hard sweeps were simulated with *θ*_A_ = 0.01, and soft sweeps were simulated with *θ*_A_ =10, as in Garud et al. 2015. The adaptive mutation was placed in the center of the chromosome. We assumed co-dominance, where a homozygous individual bearing two copies of the advantageous allele has twice the fitness advantage of a heterozygote.

To obtain a minimum of 401 SNPs for computing H12, we simulated chromosomes of length 100,000 bps for neutrality, and 350,000 bps for selection.

### Model implementations

For full details and code for model implementations, please refer to the github for this paper (https://github.com/ngarud/Harris_etal_response.git). Specifically, `documentation_Jensen_response_publication.doc` provides a README of the commands run for this paper. The script `generate_MS_commands.py` generates MS commands for all the models previously publishd. The scripts `admixture_parameters_mode_varyGrowth.py`, `admixture_parameters_mode_diffProps.py` and `admixture_parameters_mode_fixedPopSize_MS.py` generate commands for the new models tested in this paper.

The model specified in the Harris et al. supplement was coded as follows: the founding population sizes of North America and Europe were scaled by 4 *African ancestral Ne (EuroNe_anc = 10 ^log_Ne_Eur_bn^ / (4 * Ne_anc), AmerNe_anc = 10 ^log_Ne_Ame_bn^ / (4 * Ne_anc)), whereas the present day population sizes were scaled by African ancestral Ne only (scaledNeEuro = Ne_Eur / Ne_anc, scaledNeAmerica = Ne_Ame / Ne_anc), resulting in a bottleneck size that was 4 times smaller than reported by Duchen et al. 2013 (*50*) (**Figure 3I**).

### Computation of summary statistics, S, Pi, H12, and LD

S and Pi were computed from putatively neutral SNPs in short introns of the DGRP data, as described in Garud *et al.* 2015 (*34*) We used the program DaDi (*74*) to project the DGRP data down to 130 chromosomes to account for missing data. S and Pi was computed from simulations using custom python scripts.

H12 was computed from DGRP data and simulations as described in Garud et al. 2015 (*34*) using custom python scripts. LD was computed using the R^2 statistic using the same approach as described in Garud et al. 2015 (*34*) using custom python scripts. 10^7 R^2 values were averaged over to generate a smooth curve.

### Fit of a Gaussian to the distribution of H12 values

We fit a Gaussian distribution to the bulk of the distribution of H12 values by estimating the standard deviation (SD) of data within a 34.1% range of the median on either side. We ensured that data within +/−1 SD of the fitted normal was normally distributed based on a QQ plot (**Figure S4**). Since the distribution of H12 values measured from data has a long tail, we computed the number of standard deviations away from the median that the H12 value for the smallest peak was.

### Computation of Bayes Factors

We computed Bayes factors as described in Garud et al. 2015 (*34*) for two admixture models with constant population sizes in Europe and North America (**Figure 3K**). We approximated BFs using an approximate Bayesian computation approach that integrates out nuisance parameters partial frequency (PF), selection strength (s), and age of a sweep. We stated the hard and soft sweep scenarios as point hypotheses in terms of adaptive mutation rates (*θ*_A_). Specifically, BF = P(H12obs, H2obs/H1obs | soft sweep)/P(H12obs,H2obs/H1obs | hard sweep), whereby H12obs and H2obs/H1obs were computed from data, and hard and soft sweeps were simulated from a range of evolutionary scenarios.

In MSMS, when simulating selection with time-variant demographic models like the admixture model, it is only possible to condition on the time of onset of selection since the simulation runs forward in time. Thus, we assumed a uniform prior distribution of the start time of selection, ~U[0, time of admixture]. The selection coefficient and partial frequency of the sweeps were drawn from uniform priors ranging from 0 to 1.

Code and data is available for this paper at this URL: https://github.com/ngarud/Harris_etal_response.git

## Acknowledgements

We gratefully thank Pablo Duchen, Bernard Kim, Stephan Laurent, Alison Feder, Kirk Lohmueller, Daniel Schrider, and members of the Lohmueller and Petrov Labs for their generous feedback on the paper and help with implementing code.

## Supplement

**Figure S1:**
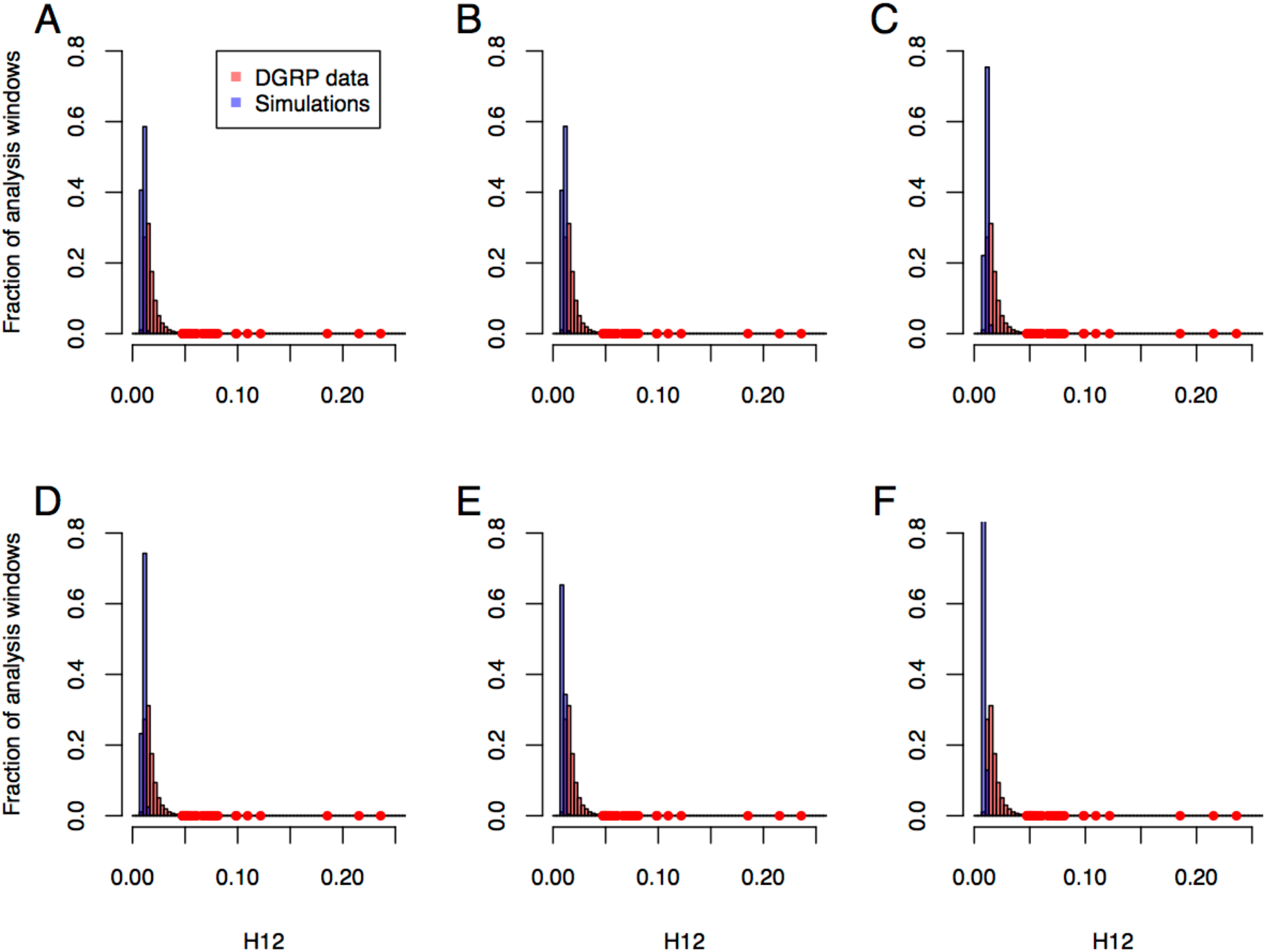
H12 values in DGRP data and in simulations of neutral demographic scenarios tested in Garud et al. 2015 (*34*). The DGRP H12 values are compared with H12 values computed in a range of simulated neutral demographic models from Garud et al. 2015. The models tested are as follows: (A) a constant *N*_*e*_ = 10^6^ model, (B) a constant *N*_*e*_ = 2.7×10^6^ model, (C) a severe short bottleneck model, (D) a shallow long bottleneck model, (E) the implemented admixture model in Garud *et al.* 2015, and (F) the implemented admixture + bottleneck model in Garud *et al.* 2015. The number of analysis windows generated for the simulated models (n=69,113) equals the number of analysis windows for the DGRP data, after excluding regions of low recombination rates. The red points indicate the H12 values for the top 50 peaks in the DGRP data.

**Figure S2:**
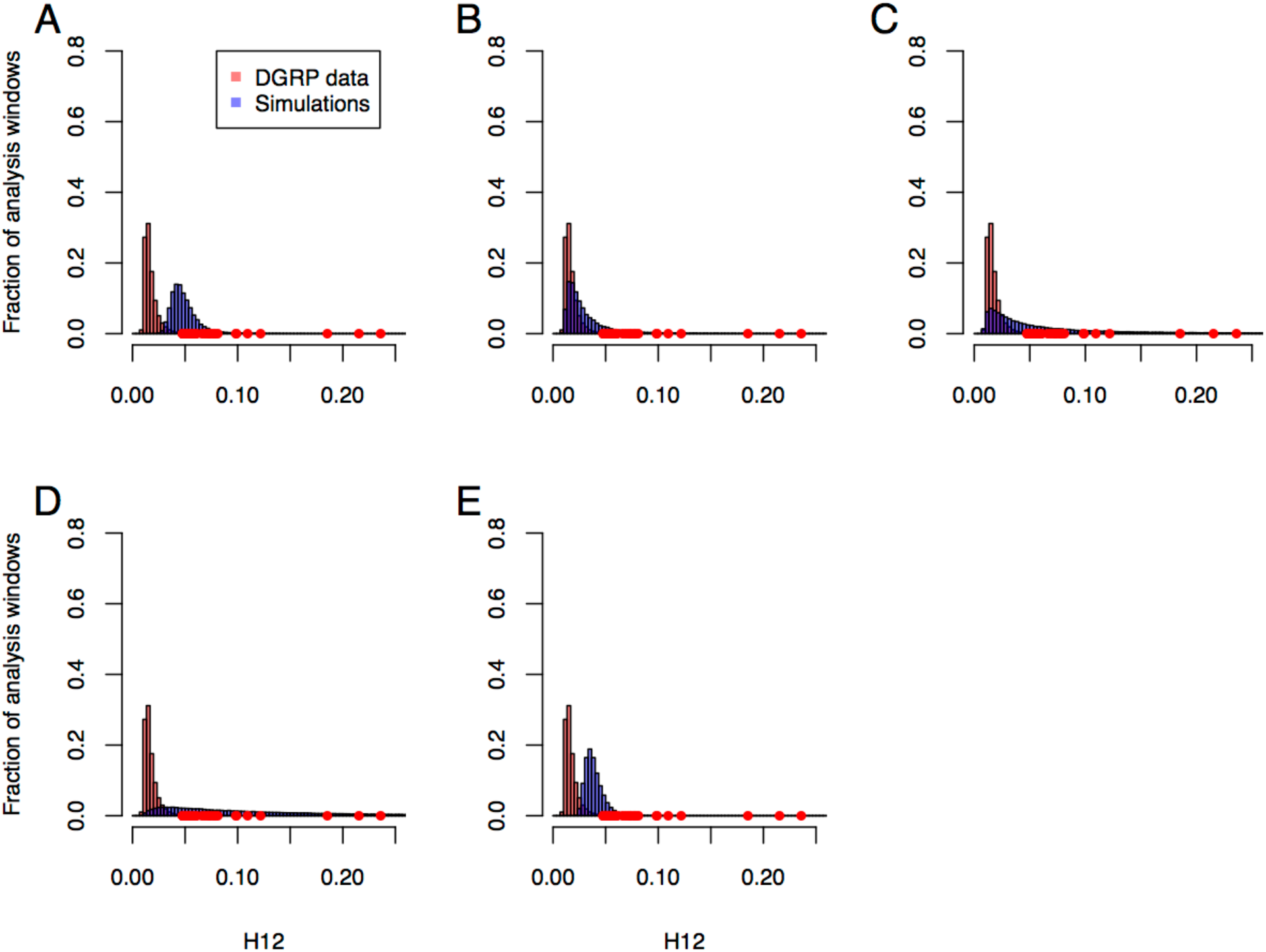
H12 values in DGRP data compared to values measured in simulations of neutral demographic scenarios from Duchen et al. (*50*), Harris et al. (*45*), and Arguello et al (*51*). The DGRP H12 values are compared with H12 values computed in a range of simulated neutral demographic models from Duchen et al. (*50*), Harris et al. (*45*), and Arguello et al. (*51*). The models tested are as follows: (A) The admixture model proposed by Duchen *et al.* 2013 (*50*), simulated with parameter values corresponding to the mode of the posterior. (B) The admixture + bottleneck model proposed by Duchen *et al.* 2013 (*50*), simulated with parameter values corresponding to the mode of the posterior. (C) The admixture model proposed by Duchen *et al.* 2013 (*50*), simulated with parameter values drawn from the posterior distribution. (D) The implemented admixture model in Harris *et al*. 2018 (*45*), simulated with parameter values drawn from the posterior distribution. (E) The admixture model proposed by Arguello *et al.* 2019 (*51*). The number of analysis windows generated for the simulated models (n=69,113) equals the number of analysis windows for the DGRP data, after excluding regions of low recombination rates. The red points indicate the H12 values for the top 50 peaks in the DGRP data.

**Figure S3:**
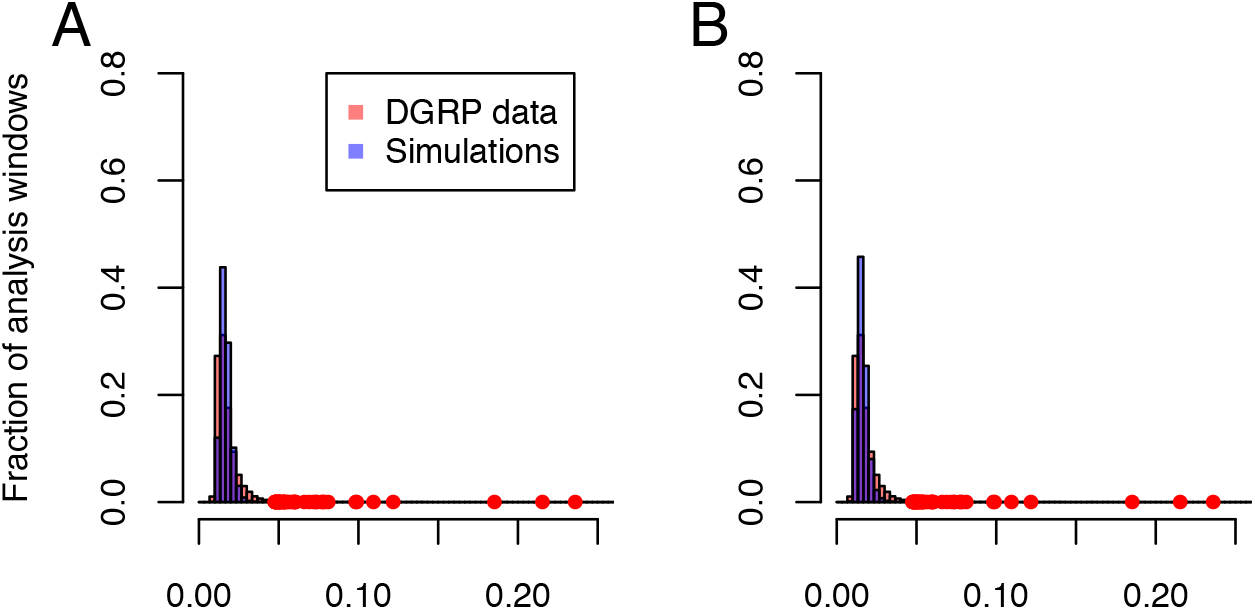
H12 values in DGRP data compared to values measured in simulations of neutral demographic scenarios inferred from this paper. The DGRP H12 values are compared with H12 values computed in two simulated neutral demographic models inferred to fit the DGRP (**Figures 3J** **and** **K**) (A) A variant of the Duchen *et al.* 2013 (*50*) admixture model where North America, Europe, and Africa have fixed population sizes. North American population size = 1.11×10^6, European population size = .7×10^6, and African population size held constant at the value inferred in Duchen et al. 2013 (*50*). (B) A variant of the Duchen *et al.* 2013 (*50*) admixture model where North America, Europe, and Africa have fixed population sizes. North American population size = 1.6×10^6, European population size = .7×10^6, and African population size held constant at the value inferred in Duchen et al. 2013 (*50*). The number of analysis windows generated for the simulated models (n=69,113) equals the number of analysis windows for the DGRP data, after excluding regions of low recombination rates. The red points indicate the H12 values for the top 50 peaks in the DGRP data.The

**Figure S4:**
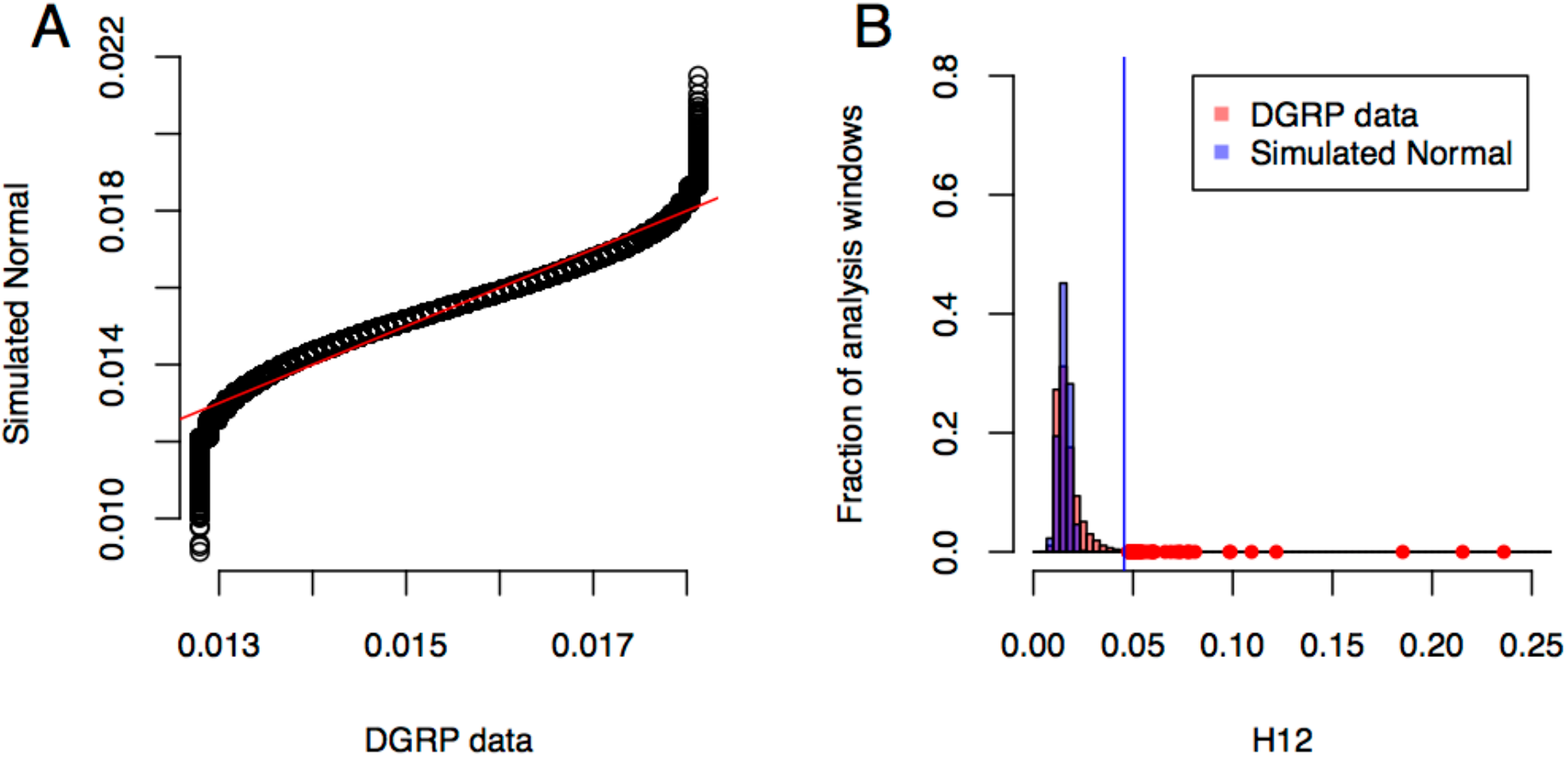
H12 values in the bulk of the DGRP data compared to a fitted Gaussian. (A) Quantile-quantile plot of H12 values within +/−1 SD of the median value in the DGRP data are compared with a random sample from the fitted Gaussian. The Gaussian was simulated with the mean equalling the median value of H12 in the DGRP data, and the standard deviation estimated from points within 1 standard deviation around the median (Methods). (B) Comparison of distribution of H12 values in DGRP data with that of a simulated Gaussian with a mean and standard deviation from (A). The vertical blue line indicates 11 standard deviations away from the mean of the simulated Gaussian distribution. The red points indicate the H12 values for the top 50 peaks in the DGRP data.

**Figure S5:**
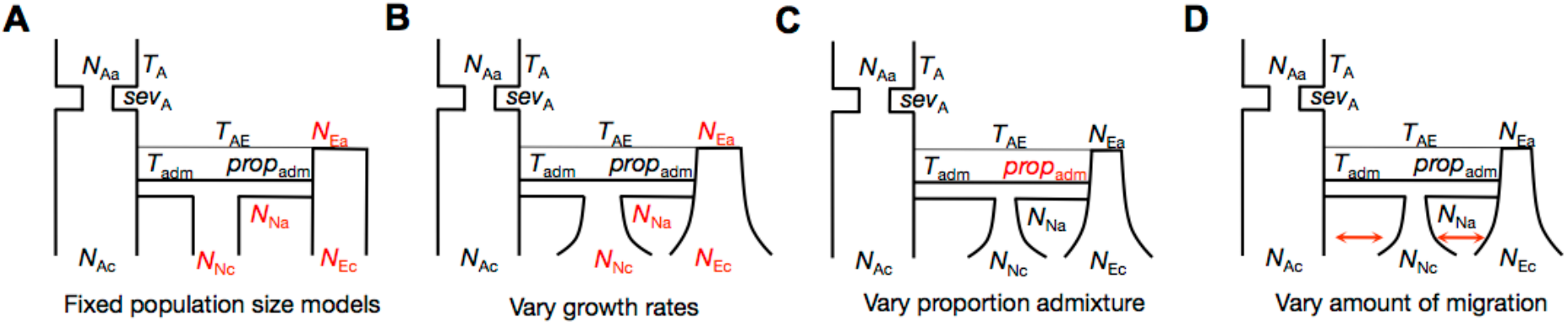
Variants of Duchen et al. 2013 (*50*) admixture models tested in this paper. We computed Pi, S, and H12 in variants of the admixture model proposed by Duchen et al. 2013 (*50*). The admixture models include: (A) constant population sizes for North America and Europe, (B) different growth rates for North America and Europe, (C) different proportions of admixture, (D) different migration rates. In all cases, the 11 parameters originally inferred by Duchen et al. 2013 (*50*) were kept constant at the mode of the parameters’ posterior distributions, unless highlighted in red. The parameters highlighted in red were varied.

**Figures S6-S10:**
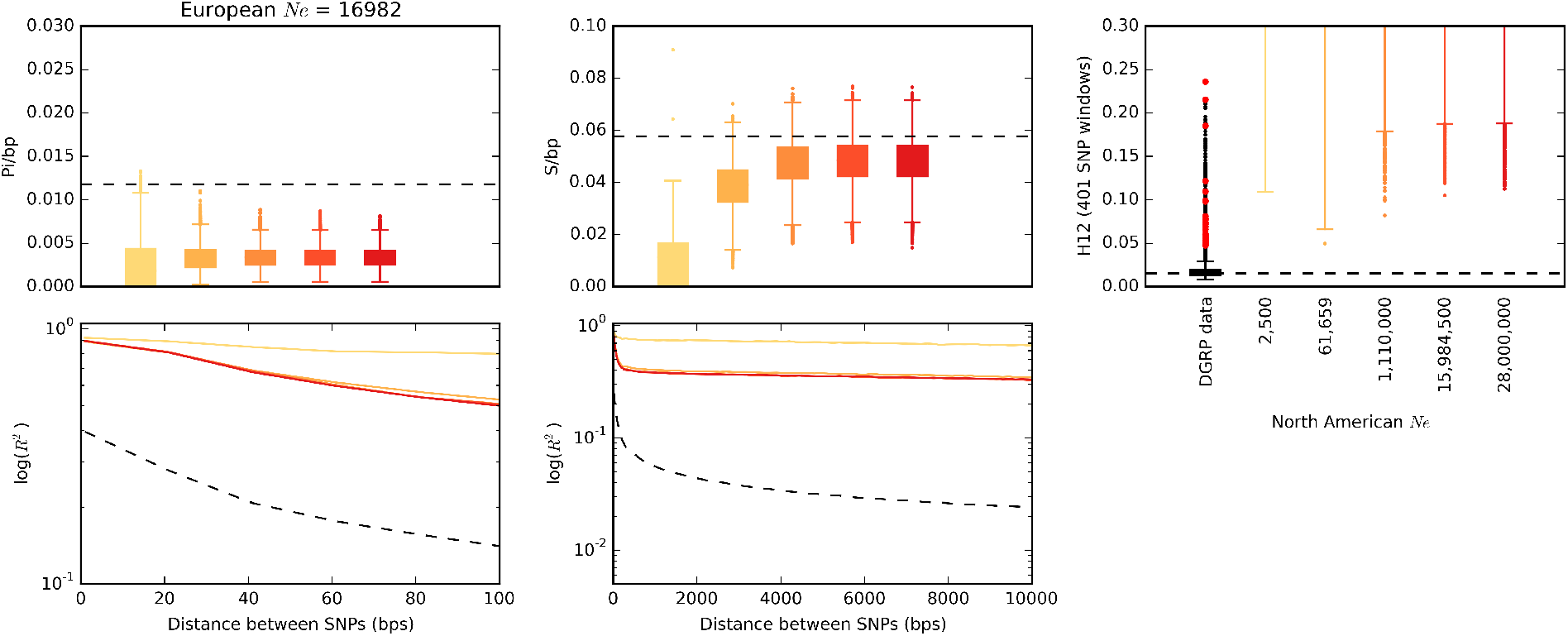
Pi, S, H12, and linkage disequilibrium measured in simulated admixture models with fixed population sizes in North America and Europe. Summary statistics S, Pi, H12, and LD were measured in admixture models with constant population sizes in Europe and North America (Figure S5A). In Figures S6 through S10, the European population size was held constant at the values 16,982, 67,608, 700,000, 2,000,000, and 9,550,000, respectively. Along the x-axis of each figure, the North American population sizes were held constant at the values 2,500, 61,659, 1,110,000, 15,984,500, and 28,800,000. These population sizes span the ranges of the 95 CI for the posterior distributions of the European and North American population sizes in Duchen et al. 2013. All other parameters in the admixture model were held constant at the mode of the posterior distribution inferred by Duchen et al. 2013 (*50*). Each boxplot is comprised of 3,000 simulations.

**Figure S7.**
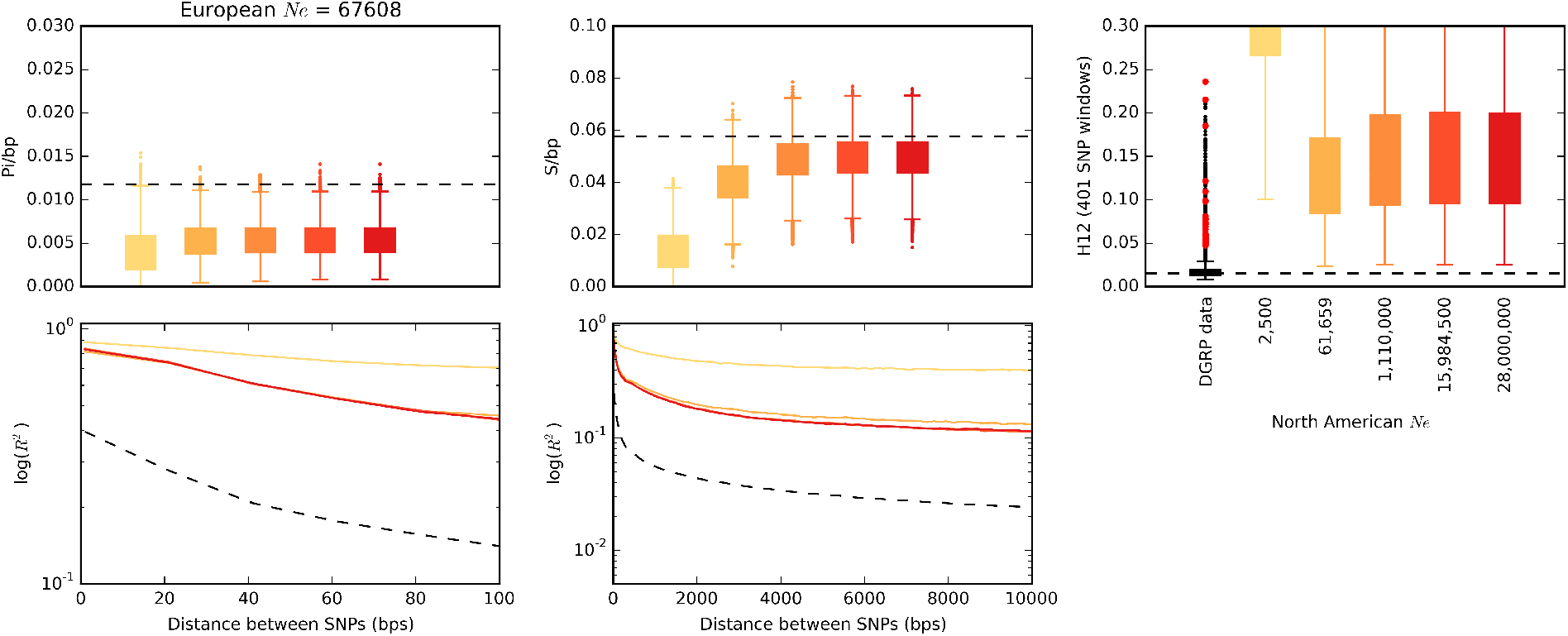

**Figure S8.**
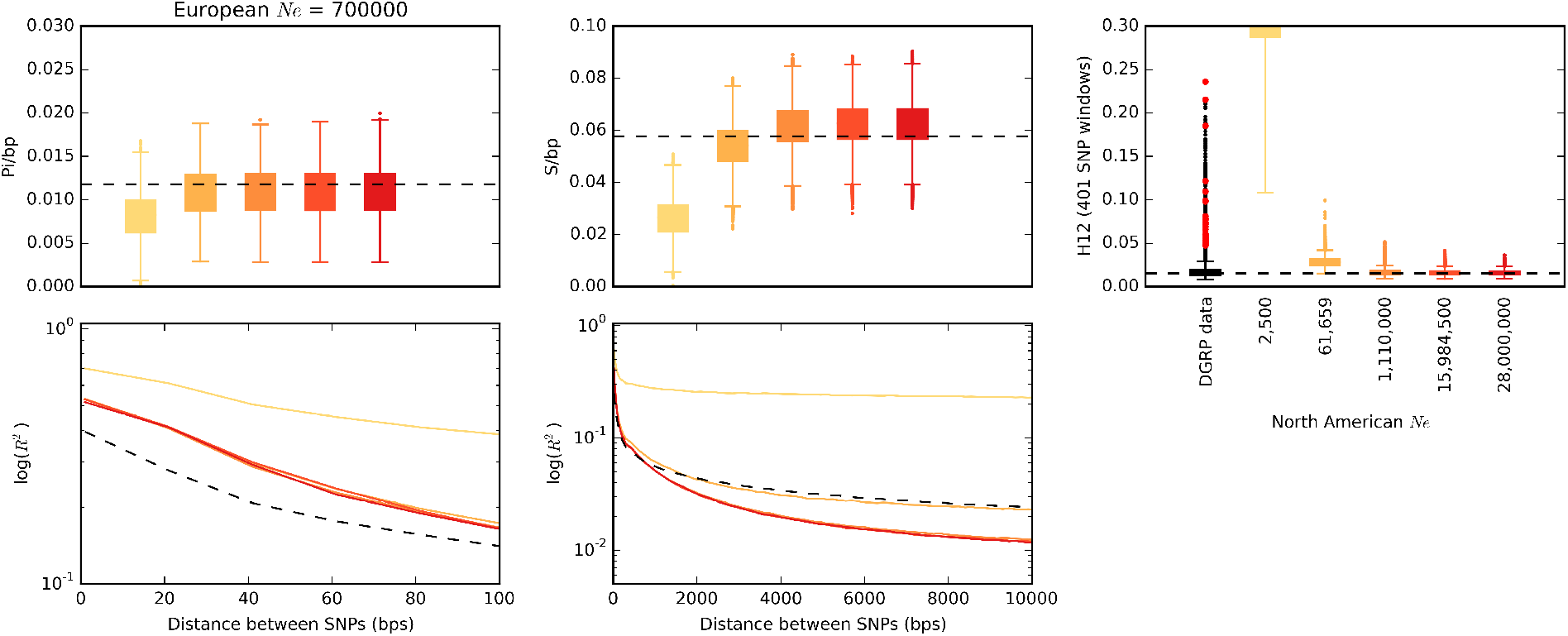

**Figure S9.**
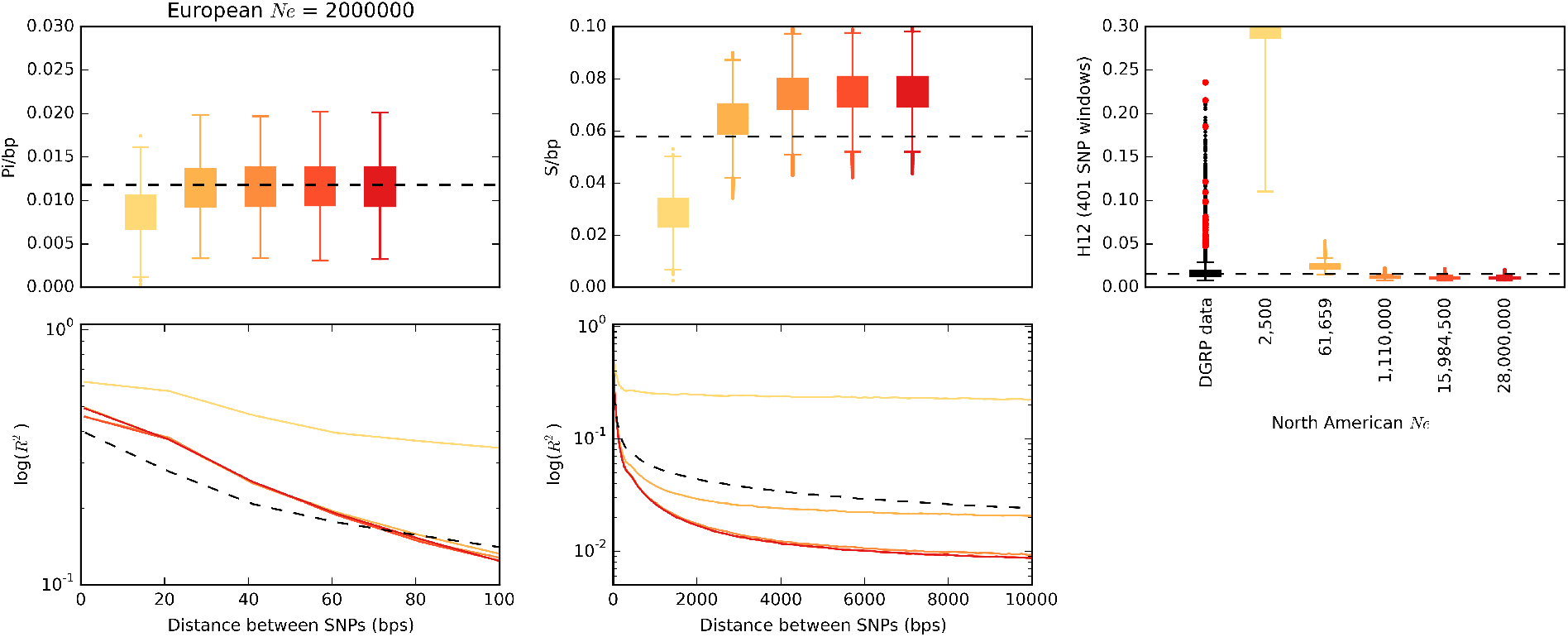

**Figure S10.**
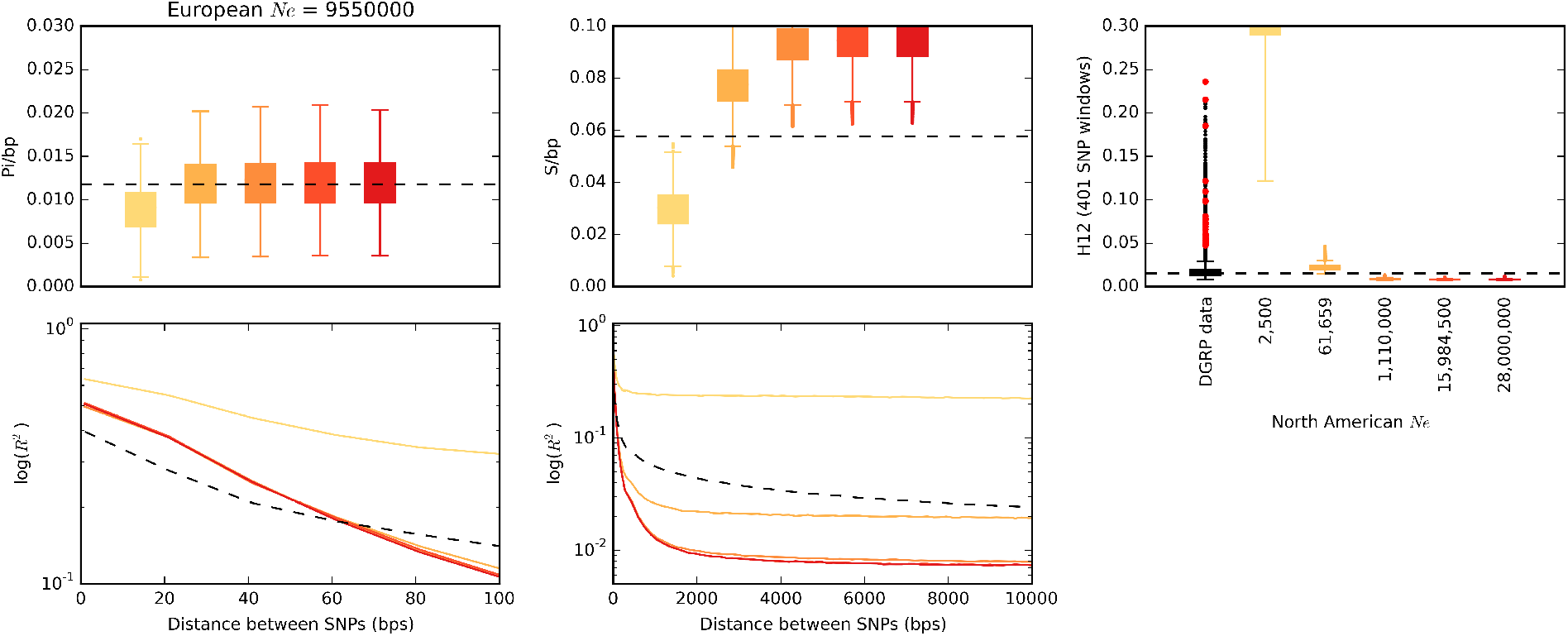

**Figures S11-S16:**
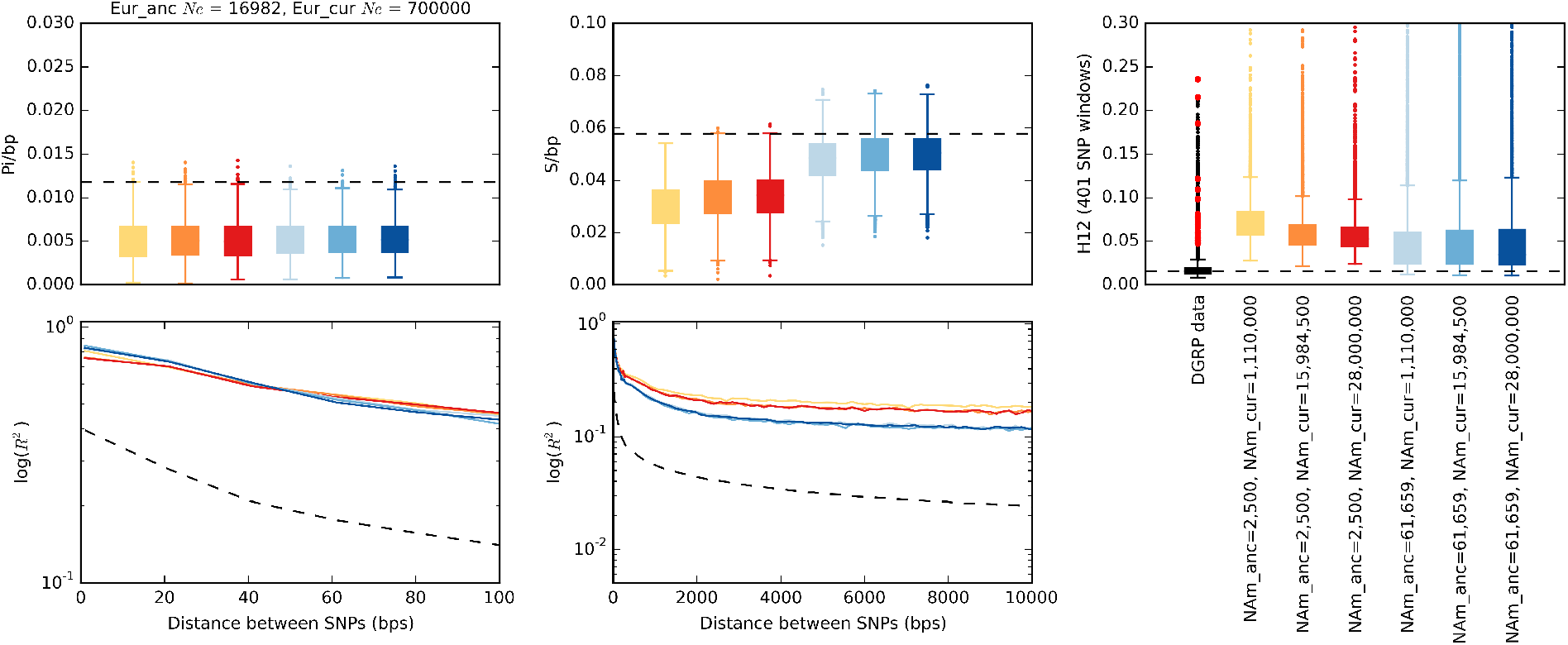
Pi, S, H12, and linkage disequilibrium measured in simulated admixture models with differing growth rates in North America and Europe. Summary statistics S, Pi, H12, and LD were measured in admixture models with varying growth rates in Europe and North America (Figure S5B). In Figures S11 through S13, the starting population size for Europe was 16,982, and ending population sizes were 700,000, 2,000,000, and 9,550,000, respectively. In Figures S14 through S16, the starting population size for Europe was 67,608, and ending population sizes were 700,000, 2,000,000, and 9,550,000, respectively. Along the x-axis of each figure, the North American starting population sizes were either 2,500 or 61,659, and ending population sizes were either 1,110,000, 15,984,500, or 28,800,000. These population sizes span the ranges of the 95 CI for the posterior distributions of the European and North American population sizes in Duchen et al. 2013 (*50*). All other parameters in the admixture model were held constant at the mode of the posterior distribution inferred by Duchen et al. 2013. Each boxplot is comprised of 3,000 simulations.

**Figure S12.**
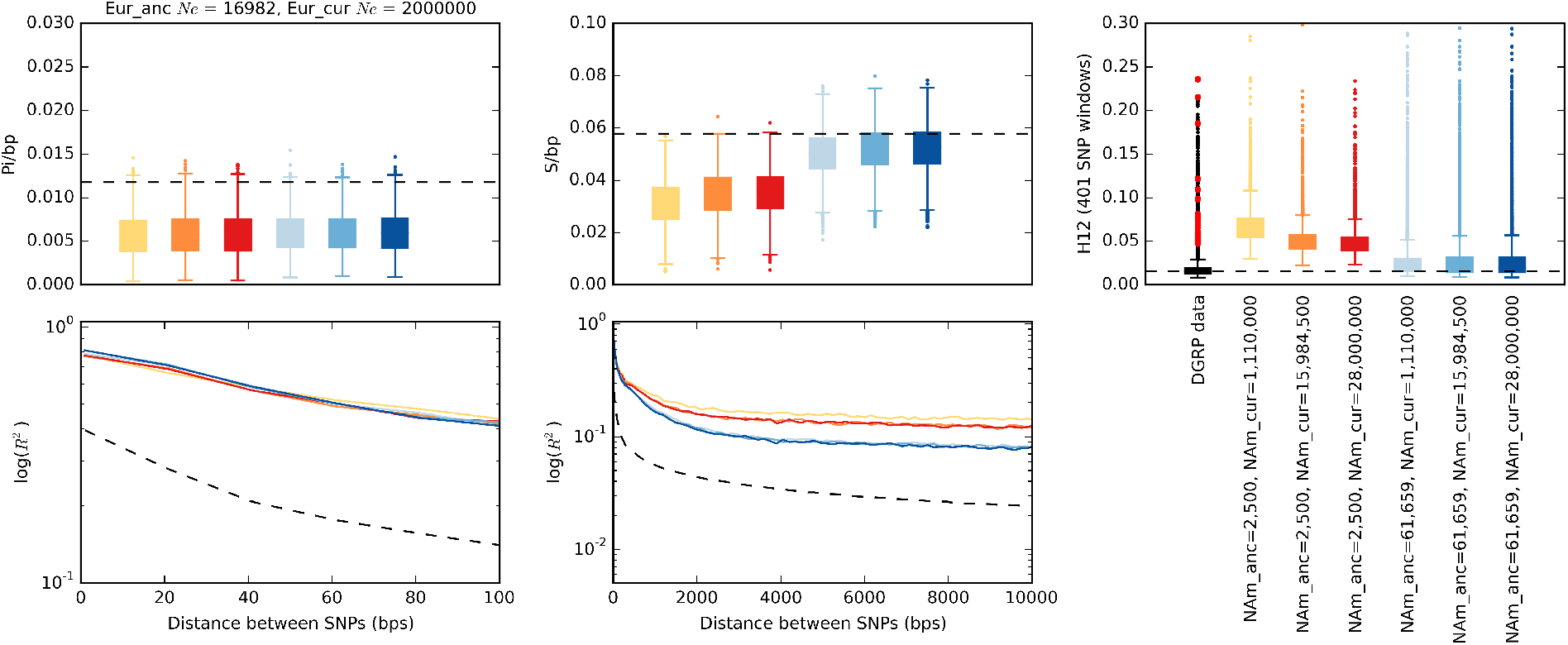

**Figure S13.**
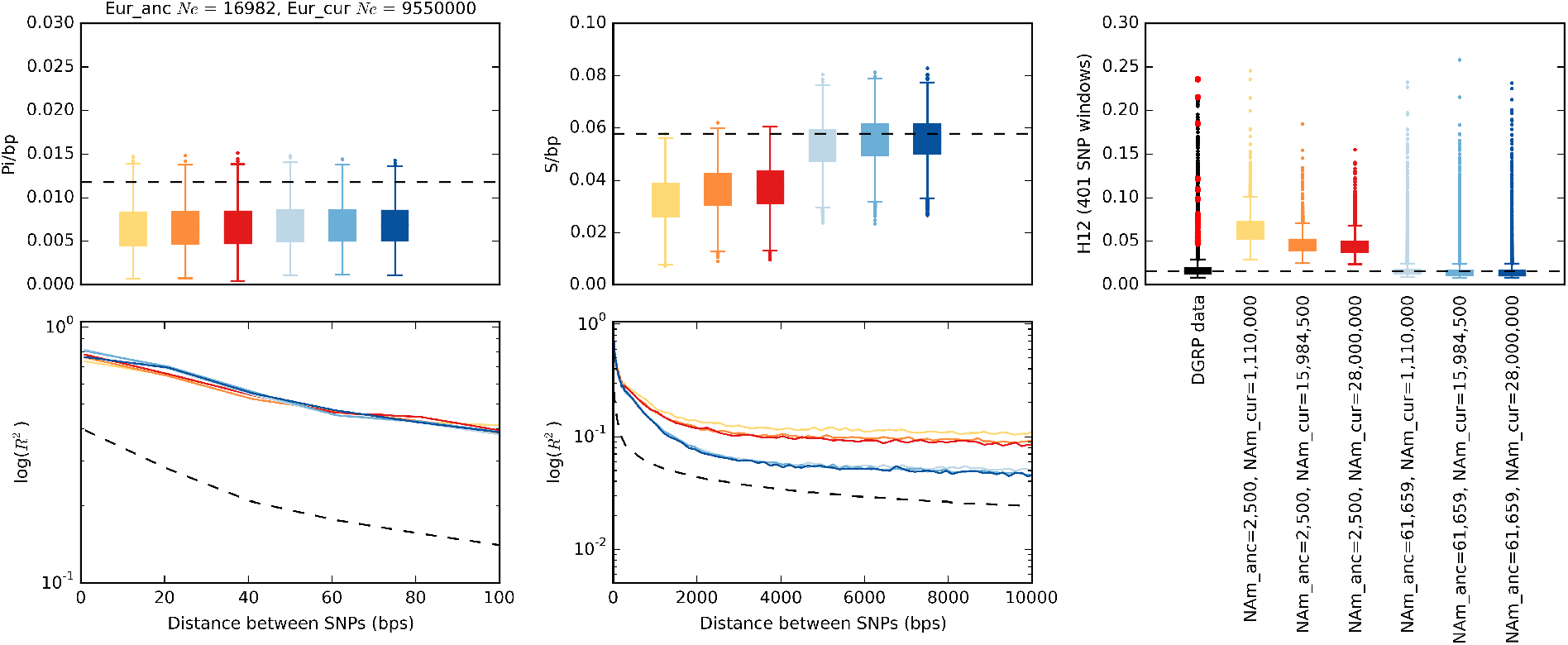

**Figure S14.**
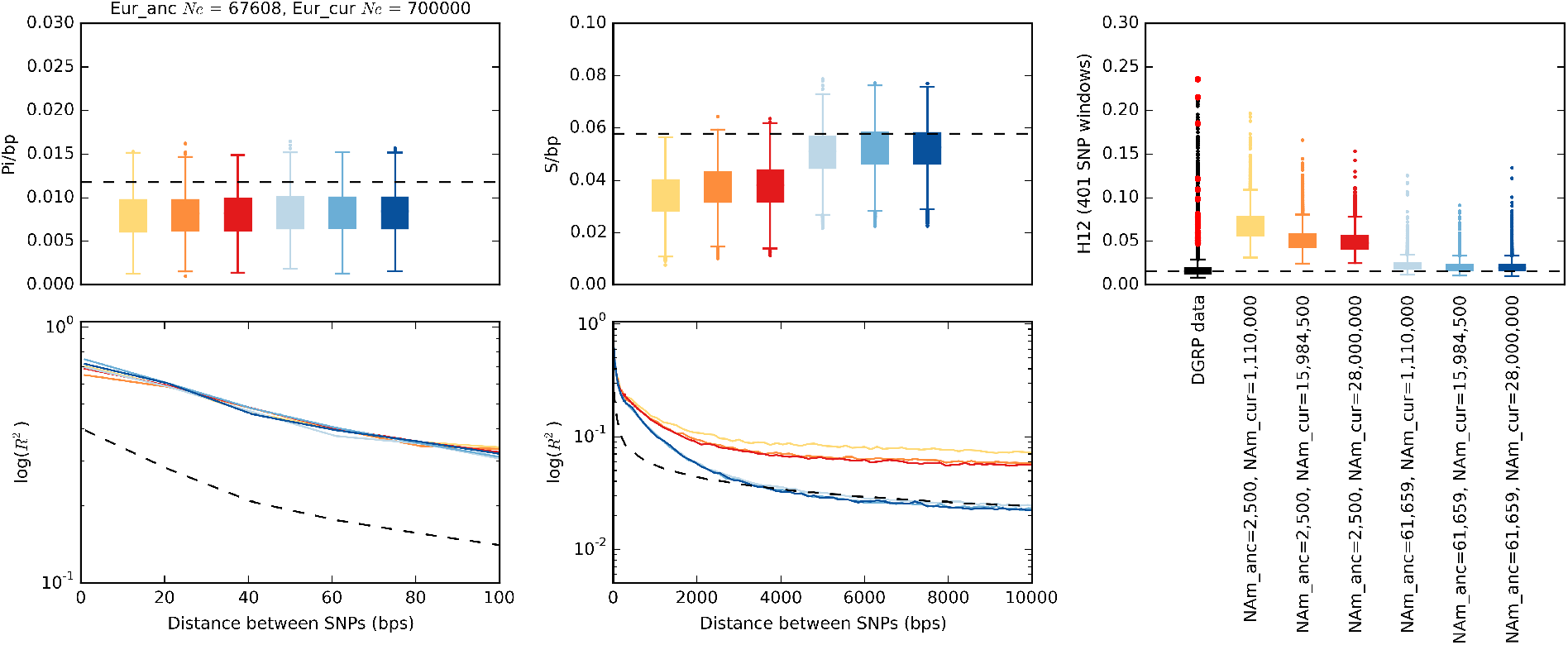

**Figure S15.**
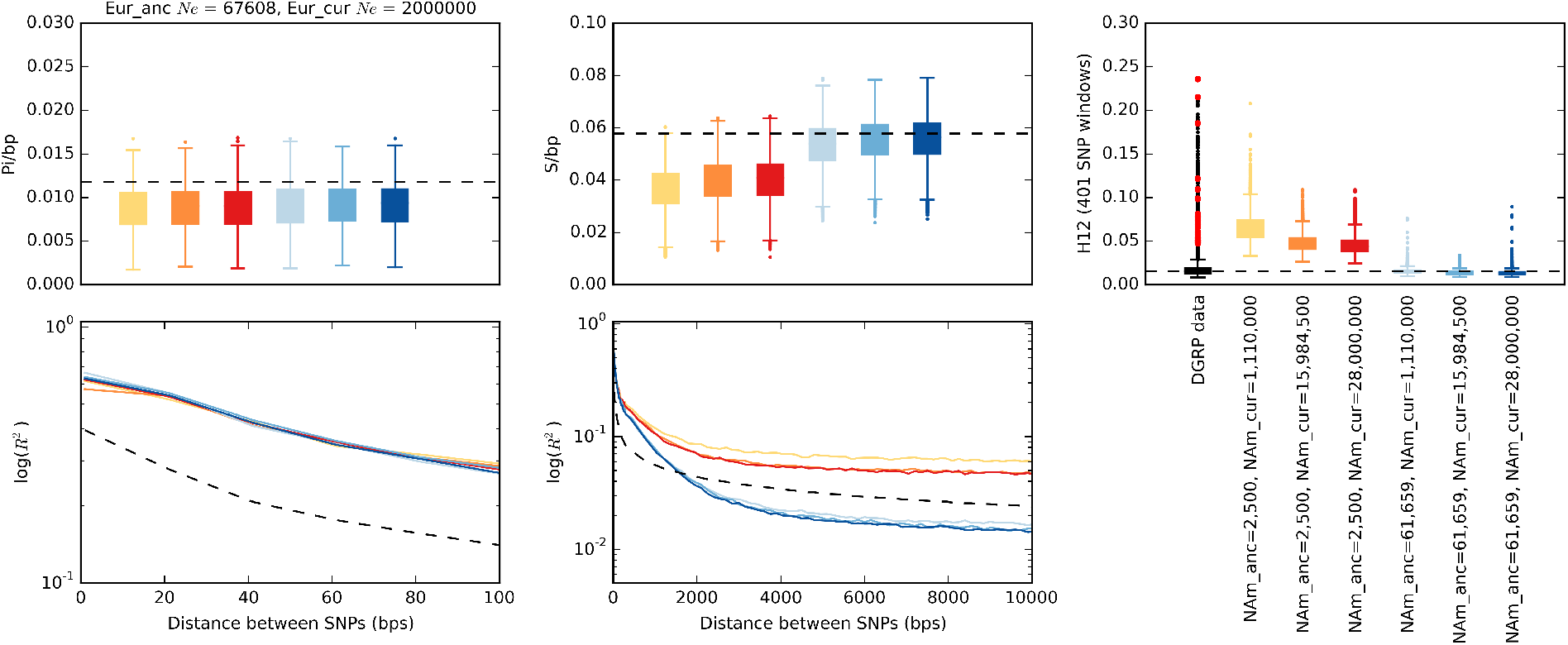

**Figure S16.**
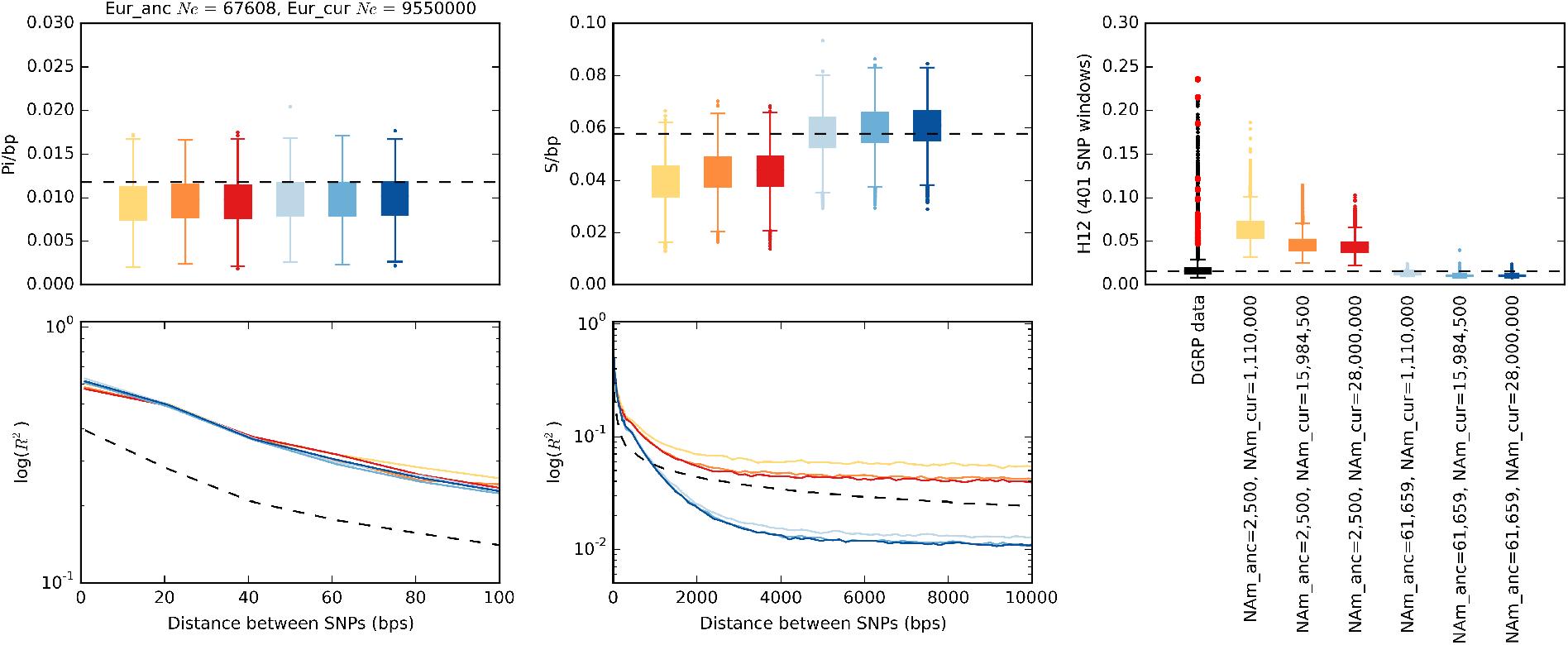

**Figure S17:**
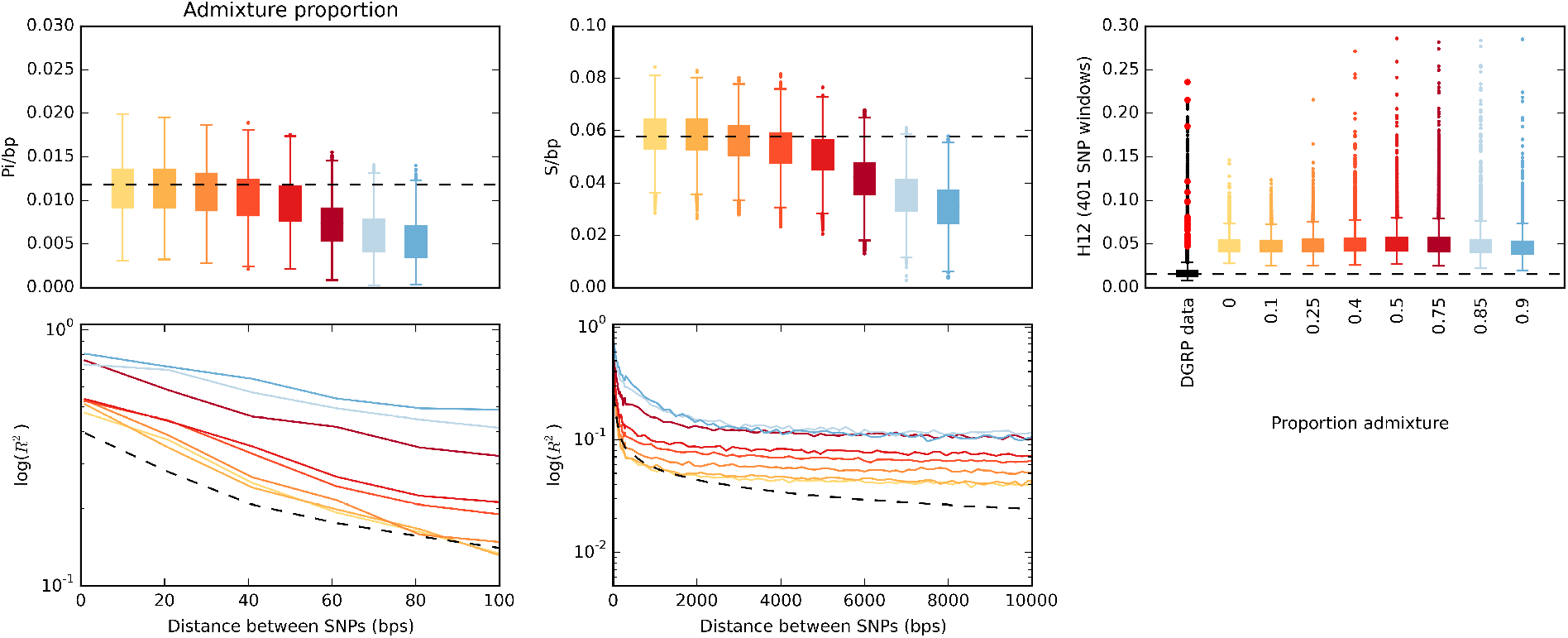
Pi, S, H12, and linkage disequilibrium measured in simulated admixture models with differing admixture proportions. Summary statistics S, Pi, H12, and LD were measured in admixture models with varying admixture proportions between Europe and North America (Figure S5C). Admixture proportions varied from 0 to 0.9. All other parameters in the admixture model were held constant at the mode of the posterior distribution inferred by Duchen et al. 2013 (*50*). Each boxplot is comprised of 3,000 simulations.

**Figure S18:**
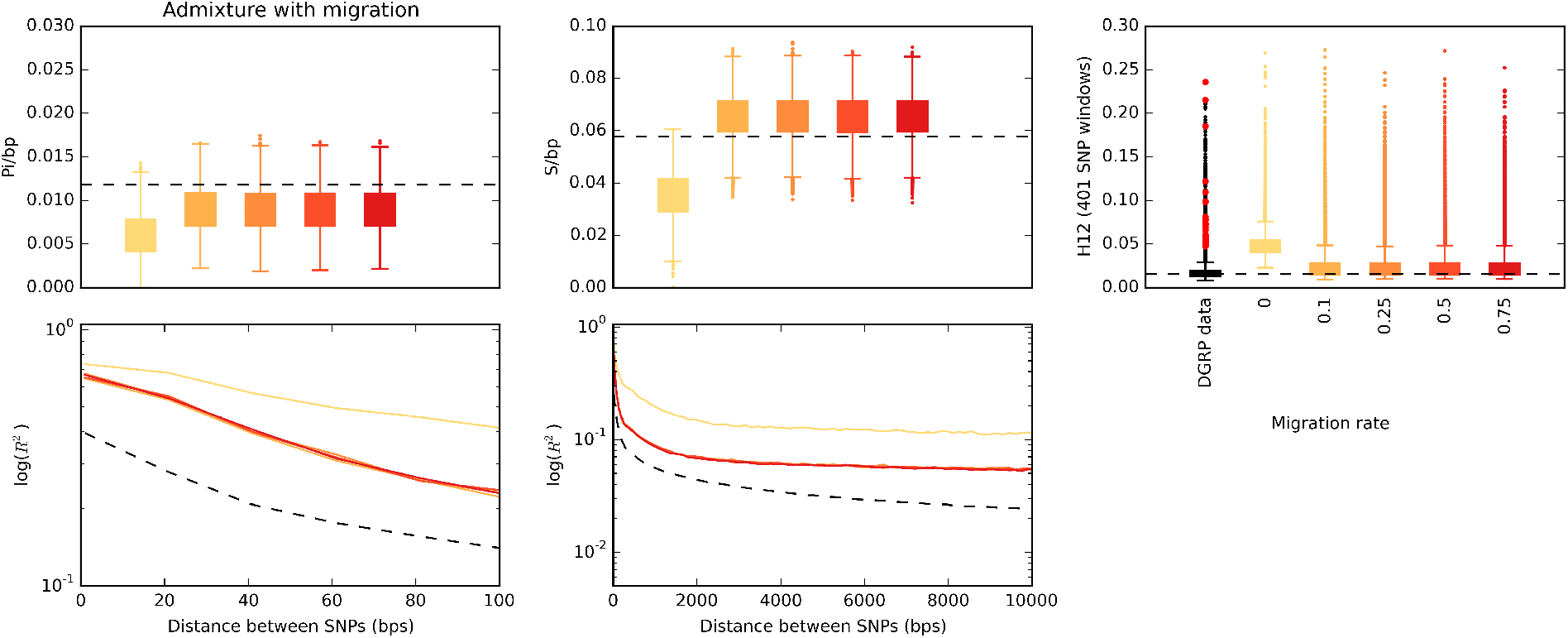
Pi, S, H12, and linkage disequilibrium measured in simulated admixture models with differing migration rates. Summary statistics S, Pi, H12, and LD were measured in admixture models with varying amounts of migration between Europe and North America (Figure S5D). Migration rates varied from 0 to 0.75. All other parameters in the admixture model were held constant at the mode of the posterior distribution inferred by Duchen et al. 2013 (*50*). Each boxplot is comprised of 3,000 simulations.

**Figure S19:**
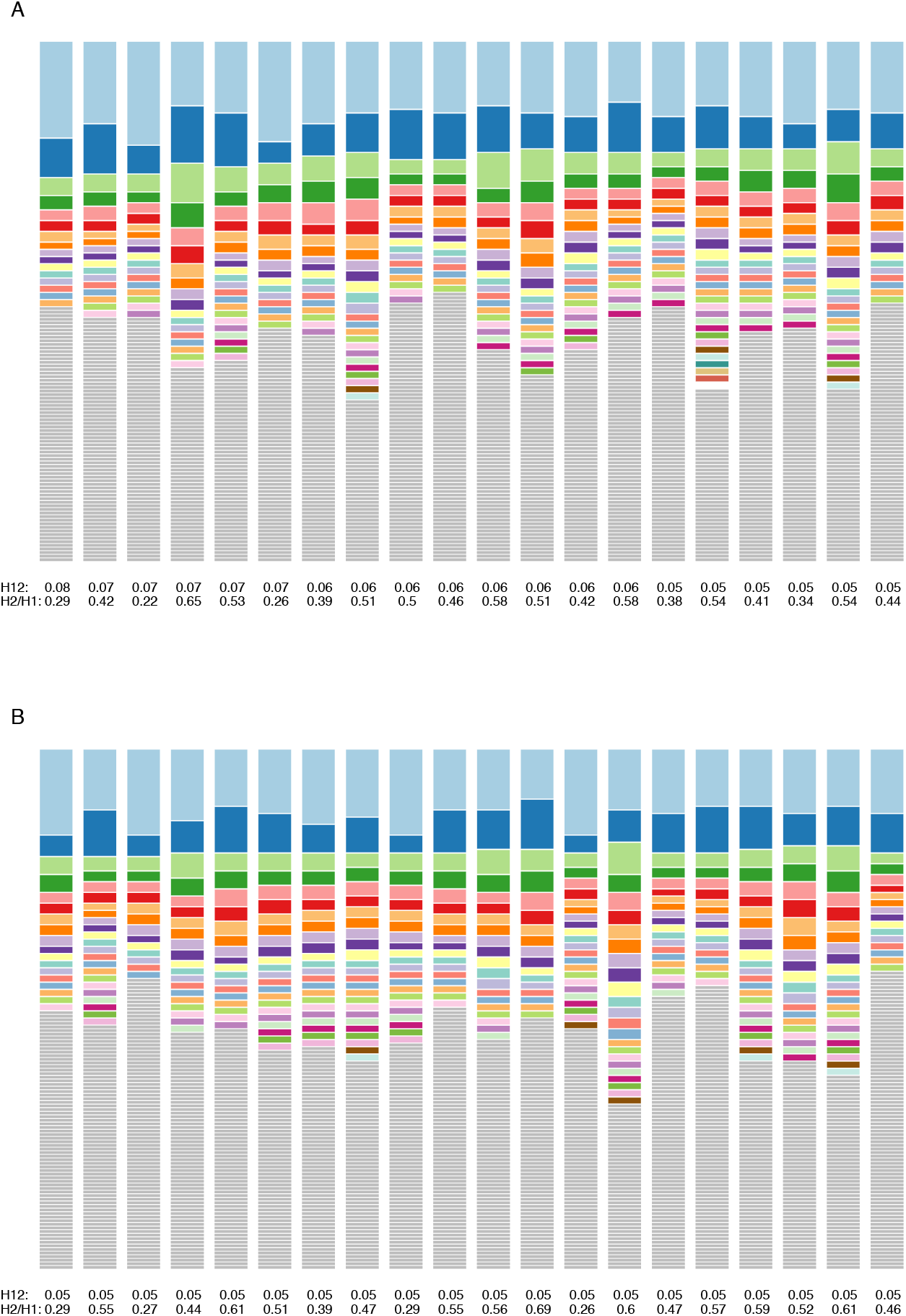
Haplotype frequency spectra for the 11^th^-50^th^ peaks. Same as Figure 7, except plotted are haplotype frequency spectra for the (A)11^th^-30^th^ and the (B) 31^st^—50^th^ peaks in the DGRP scan.

